# Thrombospondin Encapsulated Dense Particles accelerate wound healing and downregulate fibrotic phenotypes

**DOI:** 10.1101/2025.10.08.680939

**Authors:** Claire C. Staton, Marcus A. Widdess, Ashwin Jainarayanan, Anna Hoyle, Lynn Williams, Delaney Dominey-Foy, Cristina Ulivieri, Iolanda Vendrell, Roman Fischer, Cosima T. Baldari, Jagdeep Nanchahal, Dhanu Gupta, Pablo Céspedes-Donoso, Matthew J.A. Wood, Michael L. Dustin

**Author notes:** **Corresponding author** Professor Michael L. Dustin, Nuffield Department of Rheumatology and Musculoskeletal Sciences, University of Oxford. **Author Contact Information** Claire C. Staton, Marcus A. Widdess, Ashwin Jainarayanan, Anna Hoyle, Lynn Williams, Delaney Dominey-Foy, Cristina Ulivieri, Iolanda Vendrell, Roman Fischer, Cosima T. Baldari, Jagdeep Nanchahal, Dhanu Gupta, Pablo Céspedes-Donoso, Matthew J.A. Wood, Michael L. Dustin.

## Abstract

Extracellular particles (EP) released by cells are heterogenous, with extracellular vesicles (EV) believed to be the major component. To investigate different EV subpopulations, we conducted a panel screen of 15 fluorescent EV markers in 4 adherent cell lines. Nanoflow cytometry identifies TSPAN14 as the best EV marker in 3 of 4 cell lines. Surprisingly, less than 50% of EPs are labelled by the top 2 EV markers in half the cell lines. The EV marker negative EPs are detergent resistant, non-lipophilic, dense and have Thrombospondin shells by dSTORM; and were thus named Thrombospondin Encapsulated Dense particles (ThrEDs). ThrEDs are ubiquitous, contain extracellular matrix regulators and are released following cell stress. ThrEDs accelerate scratch-wound closure and downregulate the fibrotic phenotype in Dupuytren’s disease patient myofibroblasts. The TSP-1 C-terminus is exposed on the surface of ThrEDs, opening engineering applications.

## INTRODUCTION

Extracellular Particles (EP) compose an incredibly heterogeneous group of membrane and non-membrane bound particles released into the extracellular space^1^. EPs are divided into two major classes, Extracellular Vesicles (EVs) and Non-Vesicular Extracellular Particles (NVEPs)^2^. EVs are surrounded by a phospholipid membrane and function in protein turnover and as mediators of intercellular communication^3^, with three cargo compartments. The interior volume derived from cellular cytoplasm, the integral membrane glycoproteins and the extravesicular corona^4^ of attached extracellular proteins. EV populations comprise distinct sub-populations that can be separated by physical characteristics^5^.

NVEPs lack a lipid bilayer but nonetheless carry specific cargo. Exomeres^6^ and supermeres^7^ are <50 nm with distinct protein and RNA profiles. Supramolecular Attack Particles (SMAPs) are core-shell NVEPs that overlap in size with EVs with an average diameter of 120 nm^8^. SMAPs are released from multicore granules by cytotoxic lymphocytes^9^ alongside EVs with FAS ligand and interferon-γ^8^. SMAPs are built from a core of the negatively charged proteoglycan Serglycin, and positively charged cytotoxic proteins (Granzymes, INFG, PRF1) which are encapsulated by Thrombospondins (TSP1, TSP4)^10^. TSPs are extracellular matrix (ECM) proteins^11^, suggesting that SMAPs may interact with the ECM in addition to target cells.

The origins of heterogeneity reflect cell differentiation state^12^, environmental stimuli^3^, and biogenic origin^13^. Under any given set of conditions, cells will produce EPs where both EV and NVEPs contribute to biological effects but are difficult to separate. EP heterogeneity may lead to dilution of biological potency or even conflicting effects of different EP types. For example, in EV engineering, mutually exclusive loading of cargo, targeting and fusion moieties lead to a lack of efficacy^14^. Furthermore, the incorrect characterization of one EP subtype as another could cause misinterpretation of linked bioactivities. For example, a protein found in both the EV corona and an NVEP may have different functional consequences^4^.

A challenge of studying EP heterogeneity is their nanoscale often correlates with few copies of each macromolecule of interest, therefore complicating visualization^15^. Powerful fluorescence-based technologies including total internal reflection fluorescence (TIRF) microscopy^16^ and nanoflow cytometry^17^ facilitate analysis of multiple distinct features in thousands of single EPs in a systematic manner. Furthermore, super-resolution fluorescence microscopy methods like direct stochastic optical reconstruction microscopy (dSTORM) make it possible to define compartments (core-shell-corona) in sub-100 nm particles^8^. We uniformly applied these single particle and super-resolution analyses in this study to better understand heterogeneity and identify new EP types.

We initiated this study to gain insights into EV subpopulations using cell lines. A549 was used as a cancer model of EV biology based on release of small and large EVs with most characterized as exosomes based on size and morphology^18^. HeLa cells were used to study EVs derived from microvesicles budding from the plasma membrane with distinct tetraspanins and Lamp1/2 profiles^19^. Mesenchymal stem cells from cord blood (MSC(CB)) were used to evaluate 3 main subpopulations of EVs that function in tissue repair and fibrosis^20^. HEK-293T were reported to release multiple EV subtypes and because they are a standard cell line for EV production and engineering^21^. While the literature suggests these four cell types likely release exomere and supermere particles, they are not expected to release SMAPs^8^, given they are non-cytotoxic.

To better understand EP subpopulations, we focused first on EVs from HEK-293T, HeLa, A549 and MSC given their ability to release distinct subsets. We conducted an in-depth screen of 15 stably expressed putative EV markers. We found that in addition to significant cell-to-cell heterogeneity in EVs, this analysis revealed a novel population of Ca^2+^ dependent, dense glycosylated NVEPs enriched in TSPs, proteoglycans and ECM regulators. These new particles failed to be marked by any of the EV markers tested or with lipophilic dyes. These NVEP were distinct from SMAPs as they are non-cytotoxic and contained the TSP family protein cartilage oligomeric matrix protein (COMP)/TSP-5, which was not previously detected in SMAPs. Thus, we have named them Thrombospondin Encapsulated Dense Particles (ThrEDs). ThrEDs were also isolated from human plasma. ThrEDs from A549 cells and plasma were found to increase the rate of wound healing *in vitro* and reverse the fibrotic phenotype in Dupuytren’s disease derived cell cultures^22^. This novel class of NVEPs presents a new landscape for understanding the heterogeneity of EPs and may have translational potential for delivery of charged extracellular cargo that can be contained within a TSP shell.

## RESULTS

### EV marker screen highlights unexpected glycosylated NVEPs

Initially, we aimed to quantify specific EV candidate proteins released by a panel of cell lines. EV candidate proteins included tetraspanins (CD63, CD81, CD9^23^, TSPAN2 and TSPAN14^24^), Type I transmembrane proteins (PTGFRN^25^, PTTG1IP^14^ and LAMP2), Type II transmembrane protein (TFRC^26^), and cytosolic proteins (HSPA8, HSP90, SDCB, BASP1^25^ and TSG101^23^) (**Figure 1A**). First, a high throughput assay was validated in HEK-293T using transient transfection (**Figure S1A**) to determine if conditioned media (CM) yielded similar proportions of GFP^+^ EPs to size exclusion chromatography (SEC). Of 15 EV proteins only Syntenin-1 displayed a significant difference between methods (P=0.04). We concluded that the CM screening assay yielded EVs of sufficient quality to estimate differential protein association by nanoflow cytometry (**Figure S1B**). CM screening was used in pilot experiments, however final results and imaging were obtained with SEC enriched material to reduce the influence of soluble proteins.

**Figure 1.**
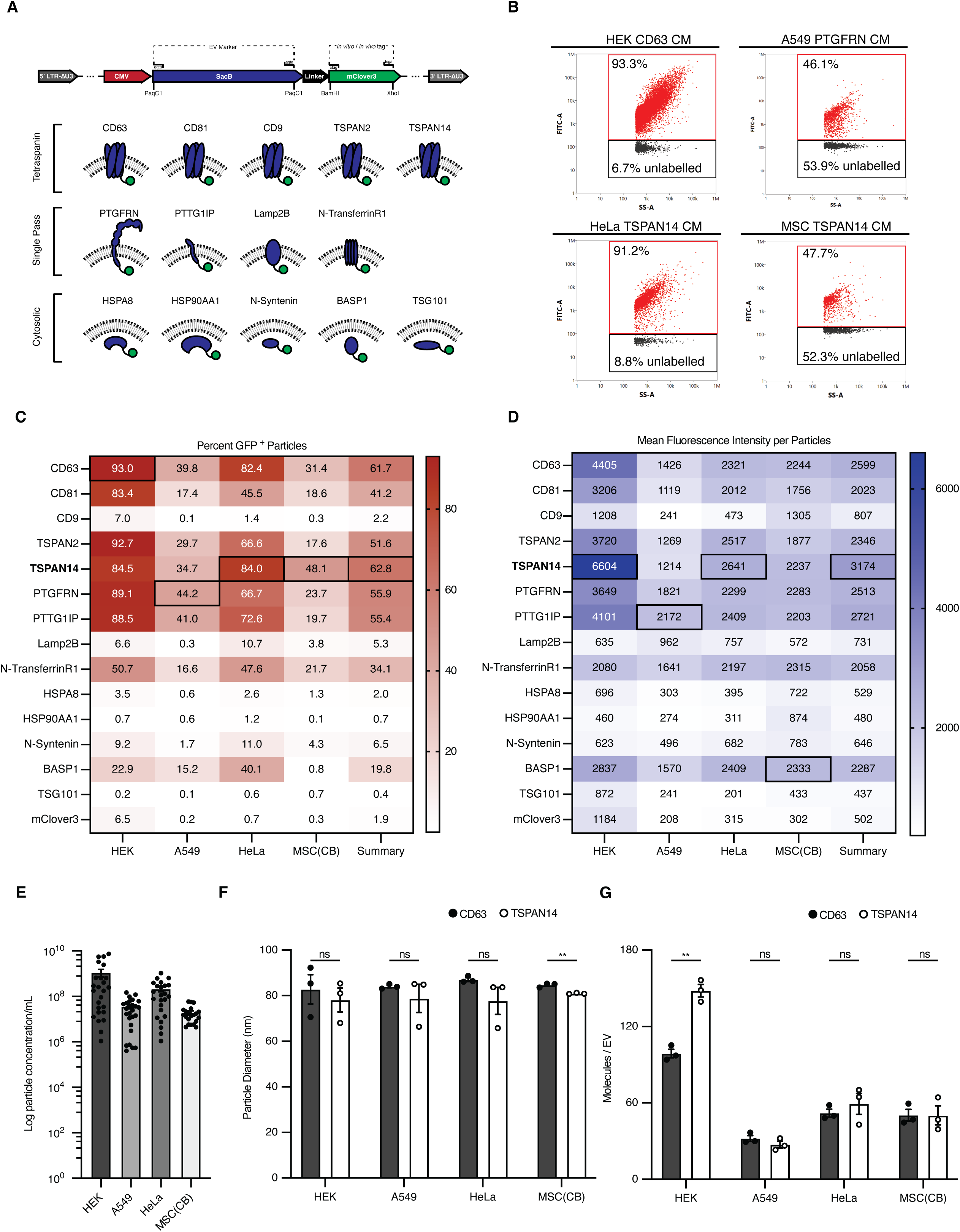
Screening putative Extracellular Vesicle markers in common cell lines. (A) Schematic depicting part of the custom lentiviral vector backbone used to clone 14 EV marker proteins classified into tetraspanin, single pass and cytosolic. Linker is the “chain”; and tag is the “ball”. (B) Nanoflow cytometry dot plot of the top EV protein marker in each cell line, showing both labelled and unlabelled populations. (C) Percentage of GFP+ (mClover3+) and (D) Mean Fluorescence Intensity (MFI) of EVs quantified by nanoFCM single-vesicle flow cytometry from conditioned media of 60 stable cell lines expressing EV marker constructs fused to mClover3. Final summary column represents the average across all 4 cell lines tested. (E) Average particle concentration measured in particles/mL via nanoFCM in HEK-293T, A549, HeLa and MSC(CB). (F) Average mean EV diameter of TSPAN14 and CD63 – mClover3 EVs measured by nanoFCM. (G) Number of molecules per EV calculated from the MFI of each EV using custom bead supported lipid bilayer (BSLB) calibration beads and MESF beads for TSPAN14 and CD63 – mClover3 EVs. Data are presented as the mean ± SEM, n = 3 biological replicates, n.s. P > 0.05, * P < 0.05, ** P < 0.01, *** P < 0.001.

To test EP heterogeneity, we stably expressed all 15 GFP fusion proteins (**Figure 1A**) in HEK-293T, A549, HeLa and MSC(CB). As expected, the EPs from each cell line exhibited different patterns of GFP expression based on both the frequency and abundance of GFP in the positive EPs. In HEK-293T, CD63 marks 93.0% of EPs (**Figure 1B, C**) and TSPAN14 had the greatest number of copies at 148 per EP (**Figure 1D, G**). In HeLa TSPAN14 labels 84.0% of EPs (**Figure 1 B, C**) with 59 copies per particle (**Figure 1G**). In A549, PTGFRN was the most frequent marker at 44.2% (**Figure 1B, C**) and PTTG1IP the most abundant at 48 per EP (**Figure 1D)**. MSC(CB) also displayed a low frequency of labelling with TSPAN14 being most frequent at 48.1% (**Figure 1B, C**) and most abundant at 50 copies per EP (**Figure 1D, G**). Cytoplasmic markers associated with EPs displayed low frequency and abundance, except for BASP1 (**Figure 1C**). Overall, TSPAN14 provides the best balance of high frequency and high abundance among all markers.

The most striking finding from this screen, however, is that A549 and MSC(CB) exhibit very low overall %EP labelling, 44.2% (PTGFRN) and 48.1% (TSPAN14), respectively (**Figure 1B, C**). To determine if low labeling proportion is due to mutually exclusive EV marker sorting^27^, EVs from cells expressing TSPAN14-GFP were stained with phycoerythrin (PE) labelled anti-CD63 to determine if there were many CD63 single positive particles that could explain the low labelling rate in some cells. While combining both TSPAN14 and CD63 increased the lower limit for the % of TSPAN^+^ EVs, the double negative EPs were 7.9% for HEK-293T, 41.7% for A549, 23.3% for HeLa, and 57.4% for MSC(CB) (**Figure 2B**). The identity of the unlabelled population became the focus of the study.

**Figure 2.**
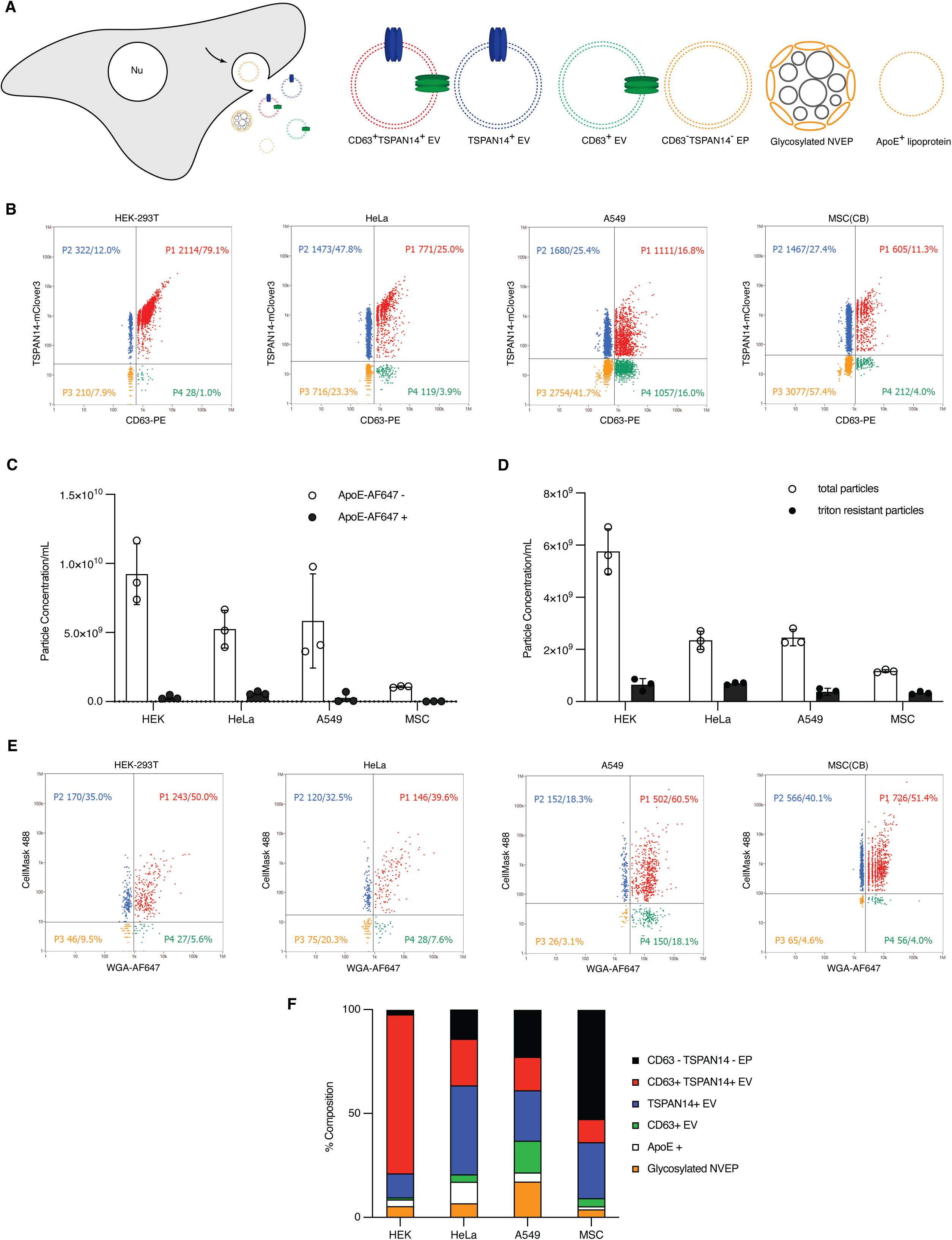
Determination of subpopulation composition using single particle flow cytometry. (A) Schematic of nanoparticle subpopulations released from cells. Particle colours are representative of quadrants in (B) (Nu = nucleus, arrow = a general representation for particle release). (B) Representative plot of single particle flow cytometry data of nanoparticle samples isolated from TSPAN14-mClover3 stable cell lines by SEC and stained with CD63-PE (P1 = top right quadrant; P2 = top left quadrant; P3 = bottom left quadrant; P4 – bottom right quadrant.) Panels (C - E) are single particle nanoflow cytometry results investigating the double negative population in (B). (C) Nanoparticle preparations from TSPAN14- mClover expressing cells isolated using SEC are probed for ApoE. Particles/mL of ApoE+ and ApoE– populations are shown. (D) Nanoparticle preparations from TSPAN14-mClover3 expressing cells isolated using SEC were measured to determine baseline particles/mL values. Samples were then treated with 0.25% Triton-X-100 and remeasured to determine the proportion of detergent resistant nanoparticles (non-membrane bound) measured in Particles/mL. (E) Nanoparticle preparations from wild type HEK-293T, A549, HeLa, and MSC(CB) cells isolated by SEC were stained with CellMask488 (membrane dye) and WGA-AF647 (glycosylated protein dye) to determine the proportion of proteinaceous particles in each cell line. Representative diagrams are shown (F) Data from (B-E) are represented as percent composition of nanoparticle subpopulations for each cell line. Data are presented as the mean ± SEM, n = 3 biological replicates.

While the double negative EPs were too large to be single lipoproteins based on light scattering^28^, we none-the-less stained for ApoE in case of aggregation. The percentage of ApoE positive vesicles was low, with HeLa having the largest proportion at 11.6% and MSC(CB) the lowest at 1.5% (**Figure 2C**). These percentages are insufficient to account for the Tetraspanin negative EPs. To assess the hypothesis that the remaining particles are lipid-based, EP preparations were isolated by SEC then treated with 0.25% Triton-X-100^29^, a non-ionic detergent that disrupts lipid vesicles, but preserves NVEPs. Triton-X-100 resistant EPs were detected in HEK-293T (5.6%), A549 (15.3%), HeLa (16.8%) and MSC (28.7%) (**Figure 2D**), confirming many of the TSPAN14^-^ CD63^-^ EP as NVEPs. Given recent findings on NVEPs, including SMAPs^8^, exomeres^6^ and supermeres^7^, whole EP preparations were stained with Cell Mask 488 (lipophilic dye) and wheat germ agglutinin (WGA)-AF647 (N-acetylglucosamine and N-acetylneuraminic acid binding lectin^30^) to classify EVs (Cell Mask 488 ^+^), non-glycosylated NVEPs (Cell Mask 488 ^−^ WGA- AF647 ^−^) or glycosylated NVEPs (Cell Mask 488 ^−^ and WGA-AF647 ^+^). This revealed the presence of glycosylated NVEPs in HEK-293T (5.6%), A549 (18.1%), HeLa (7.6%) and MSC (CB) (4.0%) (**Figure 2E**). We next performed direct stochastic optical reconstruction microscopy (dSTORM). In these experiments we used the 488 channels for anti-TSPAN14, WGA-AF568 and DiR as the lipophilic dye, which confirmed the presence of DiR^−^ WGA^+^ glycosylated NVEPs similar in size to EVs from all 4 cell lines (**Figure S2A, B**).

### Isolation of glycosylated NVEPs from A549 cells

A549 cells produced the highest proportion of glycosylated NVEPs and, therefore, we first determined the release kinetics of glycosylated NVEPs from A549 to optimize production. While the EV screen supernatants were collected after 48h, we found that we could extend the A549 culture to 96h with >80% viability and 98% confluence (**Figure S3A, B**).

We hypothesised that the glycosylated NVEP from A549 may be related to SMAPs. Given that HEK-293T and A549 do not produce Perforin1 or Granzymes, co-expressed SMAP components TSP-1 and Serglycin^8^ were investigated from day 1-5 (**Figure S3C**). The level of TSP-1 and Serglycin proteins in A549 cells was highest at day 3 and decreased significantly from day 4 to day 5. The decrease in TSP-1 and Serglycin in the cells corresponded with an increase in glycosylated NVEP (**Figure S3D**) and TSP-1^+^Serglycin^+^ particles (**Figure S3E**) on day 4 compared to day 2. Thus, glycosylated NVEP release from A549 is promoted by prolonged culture at high density, a known source of stress for A549 cells^31^, which may induce a stress responsive release of glycosylated NVEPs. Therefore, we isolated glycosylated NVEP from A549 day 4 supernatants.

Given that EVs and the glycosylated NVEPs overlap in size (**Figure 1, S2**), the Sepharose 4 Fast Flow SEC column, which has low resolution within EPs, would not separate them. We considered that TSPs are metalloproteins with multiple bound Ca^2+^ ions^32^ and the NVEP core is likely to be densely packed with proteoglycans and glycoproteins leading to high density like viral particles. Therefore, we tested 6-60% iodixanol density gradients formulated for EV-Adeno-Associated Virus (AAV) separation^33^ (**Figure 3A**). Following iodixanol gradient separation of EP from A549 cells, three EP peaks were observed (**Figure 3B**). Peak 1 (fractions 3-6) was found to be at least 68% WGA^+^CM^+^ double positive, consistent with EVs (**Figure S3G**). Peak 3 (fractions 11-12) was 91.4% WGA^+^CM^−^, consistent with glycosylated NVEPs (**Figure 3C**) and 54.7-61.0% TSP-1^+^ Serglycin^+^ (**Figure 3D, S3H**). Importantly, dSTORM images showed that the glycosylated NVEPs had a WGA^+^ and Thrombospondin^+^ shell, similar in size and morphology to SMAPs (**Figure 3E, F**). Peak 2 (fractions 7-8) contained both EVs, similar to peak 1, and glycosylated NVEPs similar to peak 3 in equal proportions (**Figure S3G**). Iodixanol gradients of A549 EPs treated with 0.1% Triton X-100 confirmed detergent resistant NVEPs in peak 2 and 3 (**Figure S3J**). TEM imaging shows EVs profiles in peak 1, NVEP profiles with a dense core, permeable to the uranyl acetate for peak 3 and both EVs and NVEP profiles in peak 2, supporting the interpretation that peak 2 results from EV- glycosylated NVEP interaction (**Figure 3G**). TSP1, enriched in peak 3 NVEPs, was detected in predominantly its full-length form, with little evidence of the 60 kda fragment (**Figure 3H**). Analytical gradients across the days of A549 culture confirm that day 4 provides the optimal balance between cell viability and the yield of the glycosylated NVEPs (**Figure S3L-N).**

**Figure 3.**
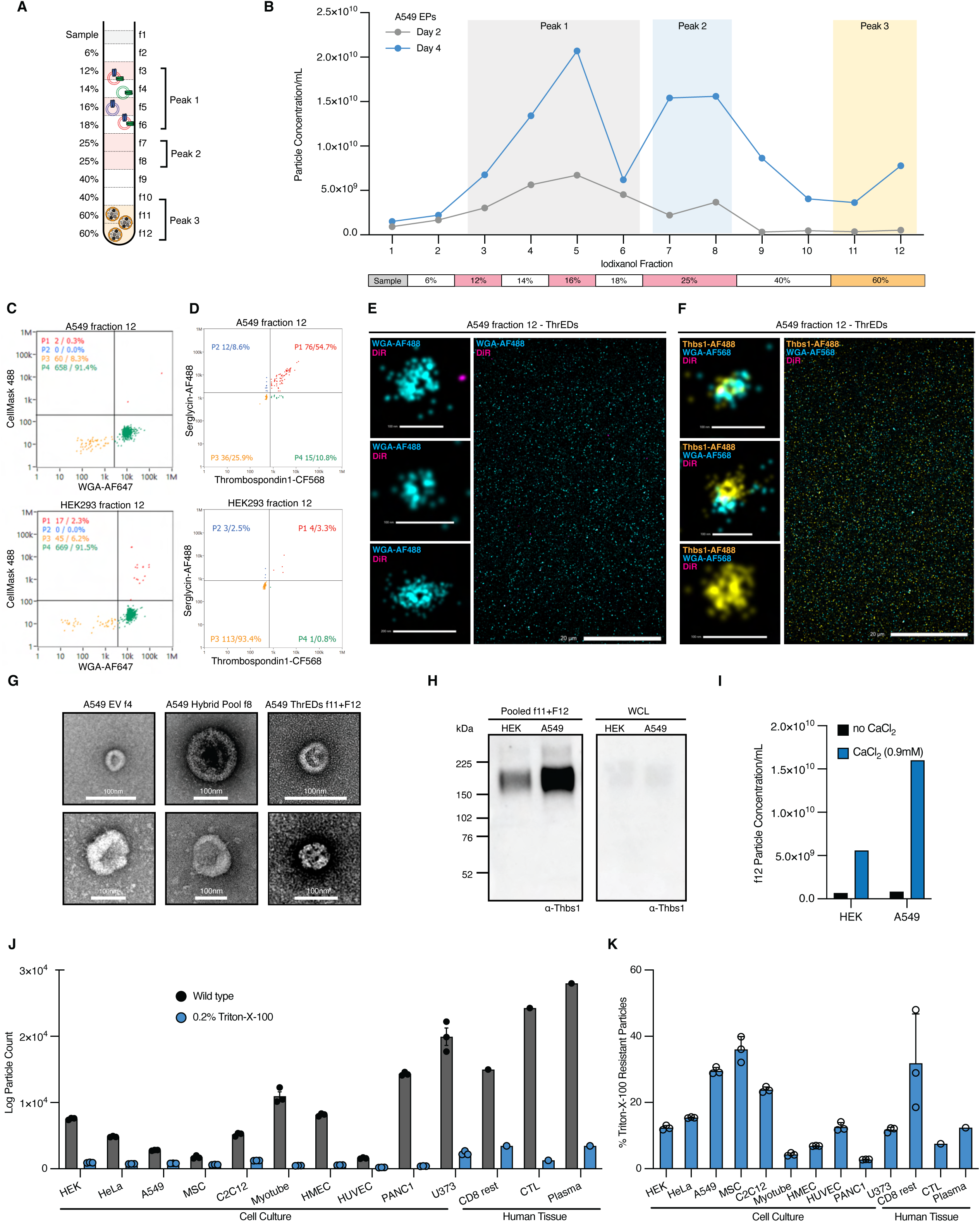
Glycosylated NVEP density-based purification and morphological characterization. (A) Schematic of iodixanol density gradient isolation protocol to generate enriched preparations of glycosylated NVEP particles for morphological and compositional analysis. Peak 1 (fractions 3-6), and Peak 2 (fractions 7-8) and Peak 3 (Fraction 11-12). (B) Migration profile of EPs, isolated at a 48 hour and 96 hour time point, following iodixanol density-based ultracentrifuge fractionation (DGUC). EV fractions are shown in the box encompassing fractions 3-6 and novel NVEP particle fractions are shown in the box encompassing fractions 11 and 12. EPs of intermediate density are shown in the box encompassing fraction 7-8. Corresponding iodixanol percentages are indicated below fraction number. (C-D) Fraction 12 particles were stained with CellMask488 and WGA- AF647 or probed for Thbs1 and Srgn for nanoflow cytometry and (E-F) DiR and WGA- AF488 or TSP1, WGA and DiR for 2D SMLM. Representative panels and images are chosen. (G) Representative negative stain electron microscopy micrographs of fractions 4, 8 and 12 are shown. (H) Western blots of pooled fraction 11+12 (glycosylated NVEPs, ThrEDs) and Whole Cell Lysates (WCL) in wild type HEK-293T and A549. ThrED particles and WCL were blotted against Thrombospondin-1. As with EVs, a loading control is not present or known. (I) The effect of PBS supplemented with calcium on glycosylated NVEP particle stability and concentration analysed using nanoflow cytometry of f12. (J-K) Nanoparticles preparations from wild type cells and human tissue isolated using SEC were measured to determine baseline particles/mL values. Samples were then treated with 0.25% Triton-X-100 and remeasured to determine the proportion of detergent resistant nanoparticles (non-membrane bound) measured in Particles/mL. Data are presented as the mean ± SEM, n = 3 biological replicates.

TSP monomers contain ∼30 binding sites for calcium ions in their C-terminal signature domain^32^. We found that scaleup of the A549 peak 3 NVEP isolations was dependent upon increasing [Ca^2+^] proportionally with input material (**Figure S4A, B**). An 8-fold increase in HEK-293T and 20-fold increase in A549 peak 3 NVEPs were obtained by maintaining 0.9 mM Ca^2+^ (**Figure 3I**). Their destabilization in divalent cation free PBS typically used in EV isolation also explains failure to previously identify peak 3 NVEPs (**Figure S4**). Ca^2+^ induced stabilization is also reversible; particles treated with EDTA largely disassembled, with only larger particles and smaller particle ‘subunits’, potentially composed of TSP1 bound to cargo, remaining (**Figure S4 C-E**). Ca^2+^ also changes particle morphology, where the shell (composed partly of TSPs) appeared bulkier and as a condensate of subunits (**Figure S4F**) that disappear in the presence of EDTA (**Figure S4G**).

Triton X-100 resistant NVEPs were identified in 6 additional cell lines; myoblasts (mouse C2C12 cell line), myotubes (differentiated from C2C12), HMEC-1 (immortalized human endothelial cells), HUVEC (primary human endothelial cells), PANC1 (pancreatic epithelioid carcinoma), and U373 (human glioblastoma). As a tissue source, we isolated them from human plasma and CD8^+^ T cells (**Figure 3J, K**).

### Glycosylated NVEPs from A549 cells and human plasma are non-cytotoxic

To investigate whether we can isolate glycosylated NVEPs from primary human tissue we collected fasted human plasma (**Figure S9A**) and compared these with glycosylated NVEPs from A549 and SMAPs from NK-92. The migration profiles of human tissue EPs were similar to A549 where 3 main peaks are observed (1, f3-4; 2, f7-8; 3, f11-12; **Figure 4A**). The particle count in f 1-2 of fasted plasma are likely circulating lipoproteins (despite fasting). The TEM analysis on these 3 peaks shows EPs like those in A549; the typical “cup” shape of EVs in peak 1, a mix of EVs and spherical dense protein particles in peak 2 and the spherical shell encapsulating the densely stained core in peak 3 (**Figure 4B**). Single particle analysis and super-resolution imaging corroborate the presence of TSPs and Serglycin in plasma ThrEDs (**Figure 4C-E, S9B**)

**Figure 4.**
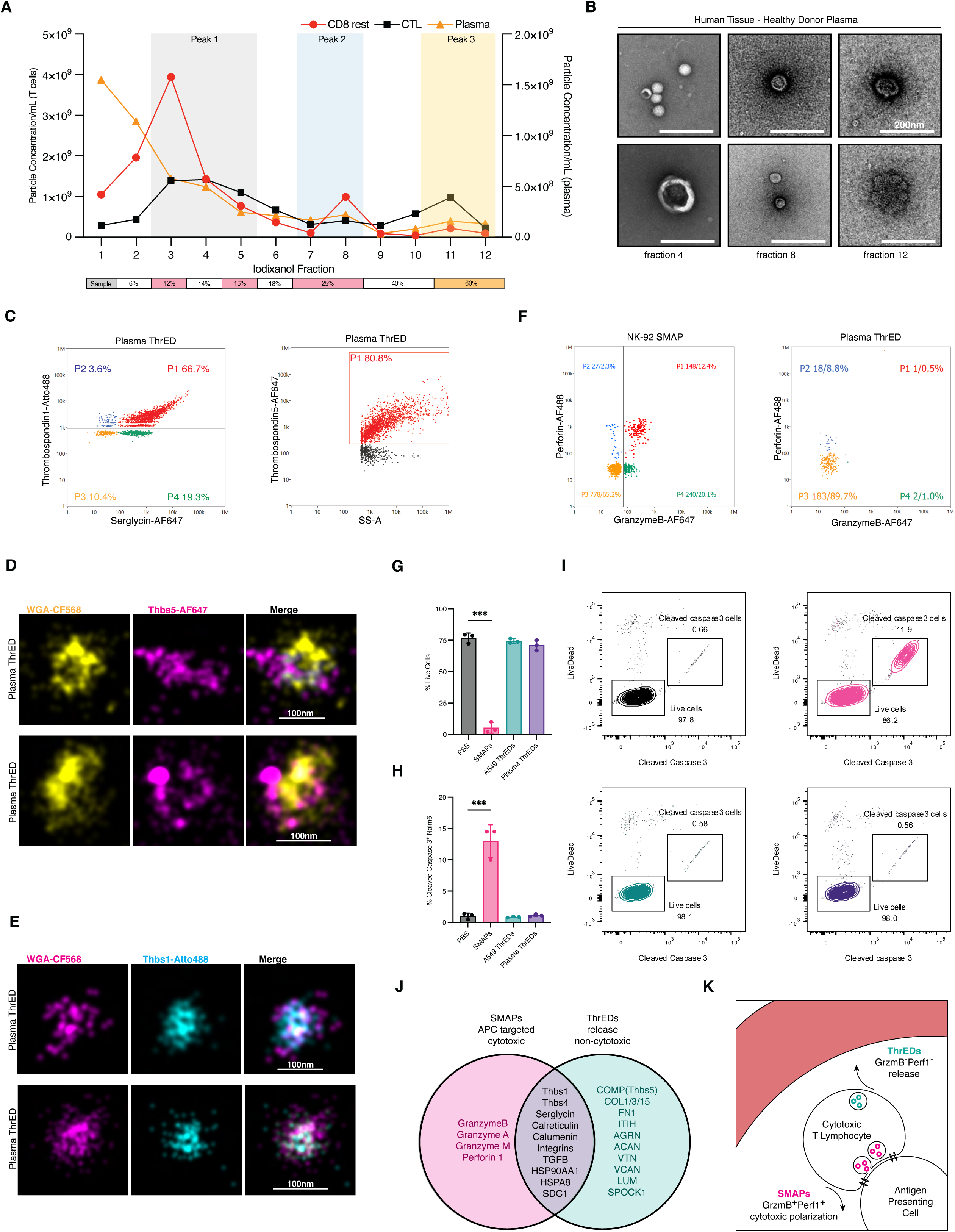
ThrEDs released from human tissue are non-cytotoxic. (A) Migration profile of EPs isolated in PBS at a 96-hour time point, following iodixanol density-based ultracentrifuge fractionation. EV fractions are shown in the box encompassing fractions 3-5 and novel NVEP particle fractions are shown in the box encompassing fractions 10-12. EPs of intermediate density are shown the box encompassing fraction 7-8. Corresponding iodixanol percentages are indicated below fraction number. (B) Representative negative stain electron microscopy micrographs of fractions 4, 8 and 12 are shown. (C) Plasma ThrEDs (f12) were probed for Thbs1, Srgn and Thbs5 (COMP) using nanoflow cytometry. (D, E) 2D SMLM dSTORM images of single ThrEDs probed for WGA (glycosylated protein dye), Thbs1 and Thbs5 (COMP) to show ring structures. (F) SMAPs from NK-92 cells and ThrEDs from plasma were probed for Granzyme B and Perforin 1 using nanoflow cytometry (cytotoxic cargo effectors). (G) Nalm6 cytotoxicity assay using SMAPs and ThrEDs to kill target cells quantified looking at cell death using a live-dead stain and Granzyme B dependent cell death by probing cleaved Caspase 3 in Nalm6 cells. (H) Representative flow plots for the quantifications in (I) are shown for each condition. (J) Venn diagram illustrating the comparative protein profiles of ThrEDs and SMAPs. (K) Schematic illustrating the difference between NVEP subtypes in CTLs. Cytotoxic SMAPs released to kill antigen presenting cells (APCs) and ThrEDs released into the extracellular space. Data are presented as the mean ± SEM, n = 3 biological replicates, n.s. P > 0.05, * P < 0.05, ** P < 0.01, *** P < 0.001.

While NK-92 cells contain both cytotoxic effectors, 98.8 – 87.3% of plasma glycosylated NVEPs are Granzyme B^−^ Perforin 1^−^, suggesting they may have a distinct function to SMAPs (**Figure 4F**). To assess this, SMAPs isolated from NK-92 cells and glycosylated NVEPs from A549 and Plasma were placed on Nalm6 cells in a killing assay. SMAPs killed with ∼98% efficiency through Granzyme B dependent Caspase 3 activation. Glycosylated NVEPs from both sources show no killing activity or activation of Caspase 3 (**Figure 4G-I**). While SMAPs and ThrEDs have similar basic proteins and morphology (**Figure 4C-F, J**), suggesting a similar overall signature, their functions may be dictated by the specialized proteins unique to each type. CTLs are particularly interesting as they release non-cytotoxic glycosylated NVEPs into the supernatant, and cytotoxic SMAPs towards the antigen presenting cell at the immune synapse (**Figure 4K, S9B, C**). Two proteinaceous particle subtypes released from one cell have diametrically opposed function depending on cellular context.

Our data suggest SMAPs are a specialized particle belonging to a more general glycosylated NVEP family similar in size to EVs but lacking cytotoxic cargo and thus are not Attack Particles. Based on the characterization so far, particles are encapsulated with a Thrombospondin shell and are denser than AAVs based on their migration into the 60% iodixanol fraction. From here on, we refer to these glycosylated NVEP as Thrombospondin Encapsulated Dense Particles (ThrEDs).

### ThrED proteomics reveal a unique marker and extracellular matrix functional signature

The absence of typical EV markers in peak 3 confirm ThrEDs are distinct from EVs. However, as with exomeres and supermeres, some EV associated proteins are present in ThrEDs including HspA8, Hsp90AA1, Syndecan-1^23^ and BASP1 (**Figure 5, S7**). Proteomics analyses conducted on EVs frequently reflect their producer cell; we hypothesized the protein profiles of ThrEDs would be variable by cell type. However, ThrEDs proteins were distinct from producer cells’ profiles (**Figure S5A, B**) and had a similar matrix composition between sources (**Figure 5A, C**), with ECM collagens, proteoglycans and glycoproteins highly enriched (**Figure S5C**). Interestingly, COMP(TSP-5), a TSP with no immune expression^34^, was abundant in samples (**Figure 5B**), making it a unique ThrEDs marker to distinguish subpopulations. Plasma ThrEDs are 80.8% COMP(TSP-5) ^+^ suggesting ThrEDs, not SMAPs, are present in circulation (**Figure 4C**).

**Figure 5.**
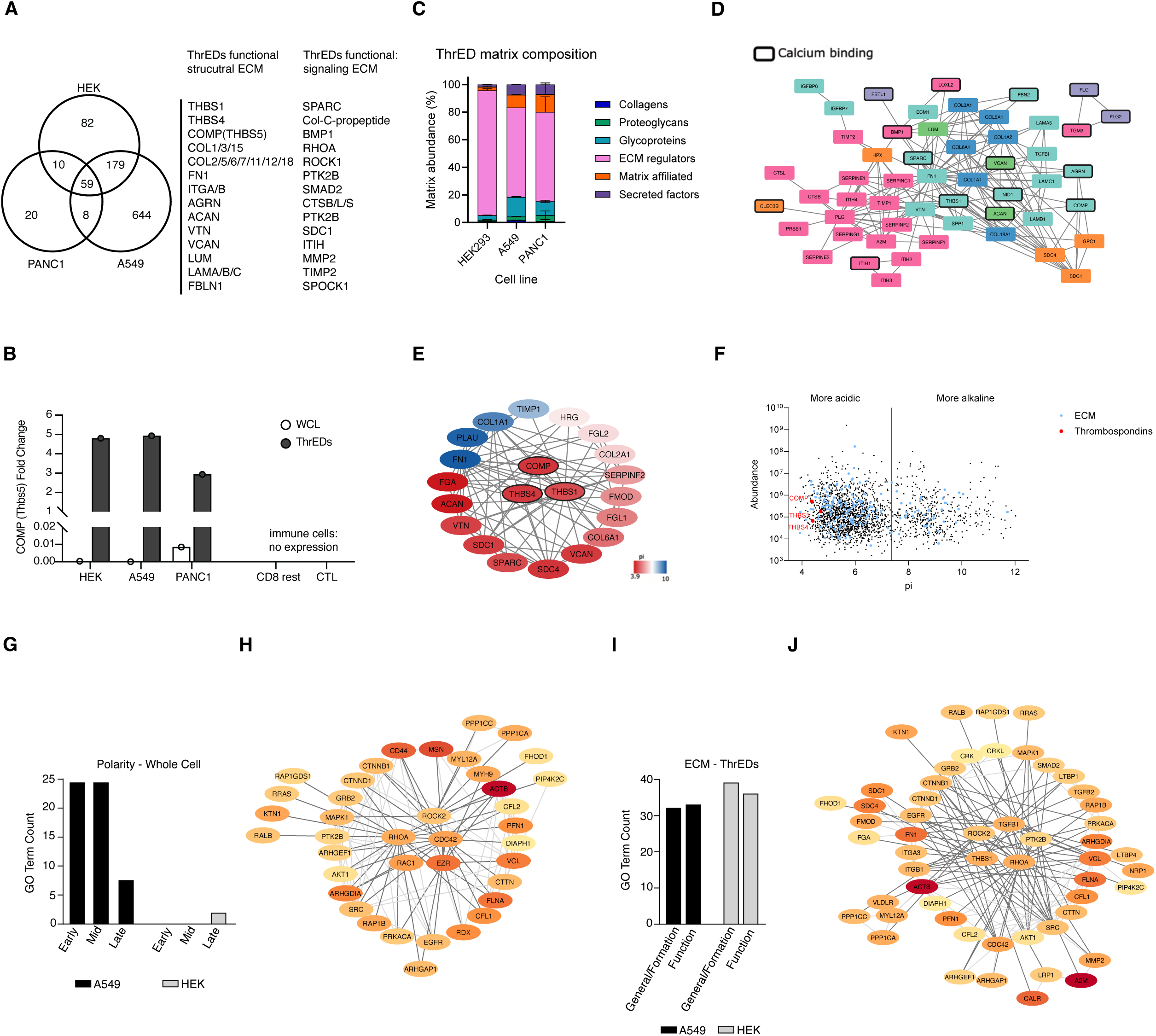
ThrED protein composition – inter/intra-particle interactions and ECM function. (A) Venn diagram of proteomics profiles in A549, HEK-293T and PANC1 highlighting common structural and signalling ECM proteins. (B) Identification of COMP (Thbs5) fold-change detected in A549, HEK, PANC1 based on detected protein abundance. (C) Overall protein matrix composition in A549, HEK-293T and PANC1 is similar and composed of collagens, proteoglycan and glycoproteins. Additional proteins are classed ECM regulators, matrix affiliated proteins (signalling, regulation, positive and negative feedback) and secreted factors. Groups were determined using MatrisomeDB. Bars from top to bottom are secreted factors matrix affiliated, ECM regulators, Glycoproteins, Proteoglycans, and Collagens. (D) Protein-protein binding interaction matrix of proteins in (C), coloured by their matrisome groups, including many calcium binding proteins. (E) Protein-protein binding interaction matrix coloured by the isoelectric point of proteins depicting charge-charge interactions between Thrombospondin shell proteins and ECM regulators identified by matrisome interacting partners, filtered for highest confidence of binding affinity (protein-protein). (F) Full protein composition of ThrEDs plotted based on isoelectric point and abundance. The red line comparatively indicates the global average protein pI. Thrombospondin shell and ECM components are largely acidic and negatively charged. (G) Number of cellular polarity related significant GO terms associated with proteins significantly upregulated in either the whole cell lysates of A549 or HEK-293T cells and (H) interaction network for the five proteins common to the greatest number of polarity GO terms. Proteins nodes are coloured by their relative abundance. Early, Mid and Late represent the different stages of initiating, establishing and maintaining cellular polarity. (I) Number of extracellular matrix related significant GO terms associated with proteins identified in A549 or HEK-293T ThrEDs and (J) interaction networks for the five proteins common to the greatest number of ECM GO terms, protein nodes are coloured by their relative abundance. Data represents n=3 biological replicates.

Thrombospondin proteins compose a part of the ThrEDs shell structure. This class of protein is particularly difficult to fold properly, given numerous cysteines (disulfide bonds), calcium dependent oligomerization, and complex glycosylation^35^. Thrombospondin accumulation at day 3 (**Figure S3C**) likely causes significant stress to the ER and protein homeostasis of the cell. Analysis of day 4 A549 lysates highlight the enrichment of proteostatic stress pathways, in particular the ER-associated Degradation (ERAD) pathway. These proteostatic proteins including SACS, SERPINF1, SERPINA2, SERPINH1, FKBP10 and CALR then localize to ThrEDs (**Figure 5, S8A-B**). Interestingly, these chaperones exhibit strong protein interactions with other abundant ThrEDs proteins including collagens to promote folding.

The isoelectric point of proteins may play a critical role in strengthening protein-protein interactions within ThrEDs. The three Thrombospondin proteins bind with high fidelity to other abundant ThrEDs proteins (**Figure 5D**), and these binding events often involve calcium or other divalent ions, which strengthens binding. Moieties with charge-charge interactions have a higher abundance in ThrEDs; Fibronectin 1 binds Thrombospondins strengthened through opposing pIs (**Figure 5E**). The overall isoelectric point profile of the ThrEDs protein matrix is primarily acidic and therefore negatively charged (**Figure 5F**). The calcium stabilization effect may be many weak interactions functioning cooperatively to maintain the ring-dense core structure as it forms a condensate.

Generally, this complement of proteins is involved in ECM dynamics, cell migration and wound healing (**Figure S6)**. ITIH1, 2, 3 and 4, which bind hyaluronan (HA; **Figure 5D, S5D**) bind water to hydrate the ECM in addition to serving as a scaffold and signaling molecule^36^. Collagen 1A, the most abundant form of collagen^37^, is another universally abundant ThrED component. SERPINE1, SERPINI1, SERPINB3, SDC1, FSLT1, IGFBP6 (**Figure 5D, S5**) are all involved in ECM modification to facilitate cell migration. In addition, SERPINF2, FLG2, PTPRK, CLSTIN1, CNTN1, CSTA, NRCAM and DSC1, common to samples (**Figure 5A)**, initiate the cell’s orientation and establish polarized function through adhesion functionality.

To understand why A549 produce more ThrEDs than other cell lines, we analysed their cell lysates at day 4. We hypothesized the strong polarization in CTLs was important for the release of glycosylated NVEPs. The analysis of differentially expressed proteins found 2 cell polarity GO terms in HEK293, while A549 had 53 involved in initiation, establishment, and maintenance of polarization (**Figure 5G, S6D**). Many of the polarity proteins then localize into released ThrEDs and sort into the ECM functional matrix web (**Figure 5H, I, J).** Furthermore, despite HEK293’s few lysate polarity proteins, similar proteins localize abundantly into HEK293 and A549 ThrEDs including RHOA, ROCK2, CDC42, MAPK1, AKT1 (**Figure 5H-J**). A mechanism to sort components into ThrEDs regardless of copies per cell may exist, accounting for the similarity in overall protein matrix composition (**Figure 5C**).

### ThrEDs from human tissue and cell culture function in wound healing and fibrosis reversal

The extracellular matrix is a network of proteins and polysaccharides surrounding cells, providing dynamic support to regulate cellular function. ECM remodeling is tightly controlled to maintain homeostasis. Deregulation causes pathologies including fibrosis, defined by excessive ECM deposition^38^. Dupuytren’s disease is characterized by excessive local accumulation of myofibroblasts and ECM within palmar fascia nodules leading to permanent finger flexion deformities^22^.

A wounding assay monitoring cell migration across a uniform scratch was used to investigate ThrEDs mediated wound healing. ThrEDs from A549 and plasma both induce significantly faster wound closure, in a dose dependent manner (**Figure S10A, B**), due to the activation of Rac and phospho-focal adhesion kinase (p-FAK) in recipient A549 cells (**Figure S10D, E**). ThrEDs induce significant actin cytoskeleton reorganization, not only at the migrating edge, but the entire wound, likely contributing to closure within ∼24h compared to ∼72-120h for PBS (**Figure S10B, C**).

Primary myofibroblasts isolated from Dupuytren’s disease (DD) patient nodules were then investigated. Fibroblasts migrate in the early wound secreting ECM and signaling molecules. Myofibroblasts are activated fibroblasts with reduced migration capacity which contract and remodel the ECM^22^. DD myofibroblasts treated with ThrEDs and hybrids significantly increase wound closure (**Figure 6 A-C**) through increased Rac, decreased p-FAK and reversion to spindle morphology (**Figure 6D-F**). Conversely, EVs cause myofibroblasts to migrate more slowly by increasing a contractile myofibroblast phenotype. Interestingly, ThrEDs may also function extracellularly. ThrED proteins Fibronectin 1, Collagen1A (**Figure 6I**) and Actin have significantly greater puncta, 500nm or less, in the scratch wound area compared to control (**Figure 6G, H**). ECM synthesis requires a nucleation point^39^, which may be facilitated through ThrED deposition in the wound area^40^.

**Figure 6.**
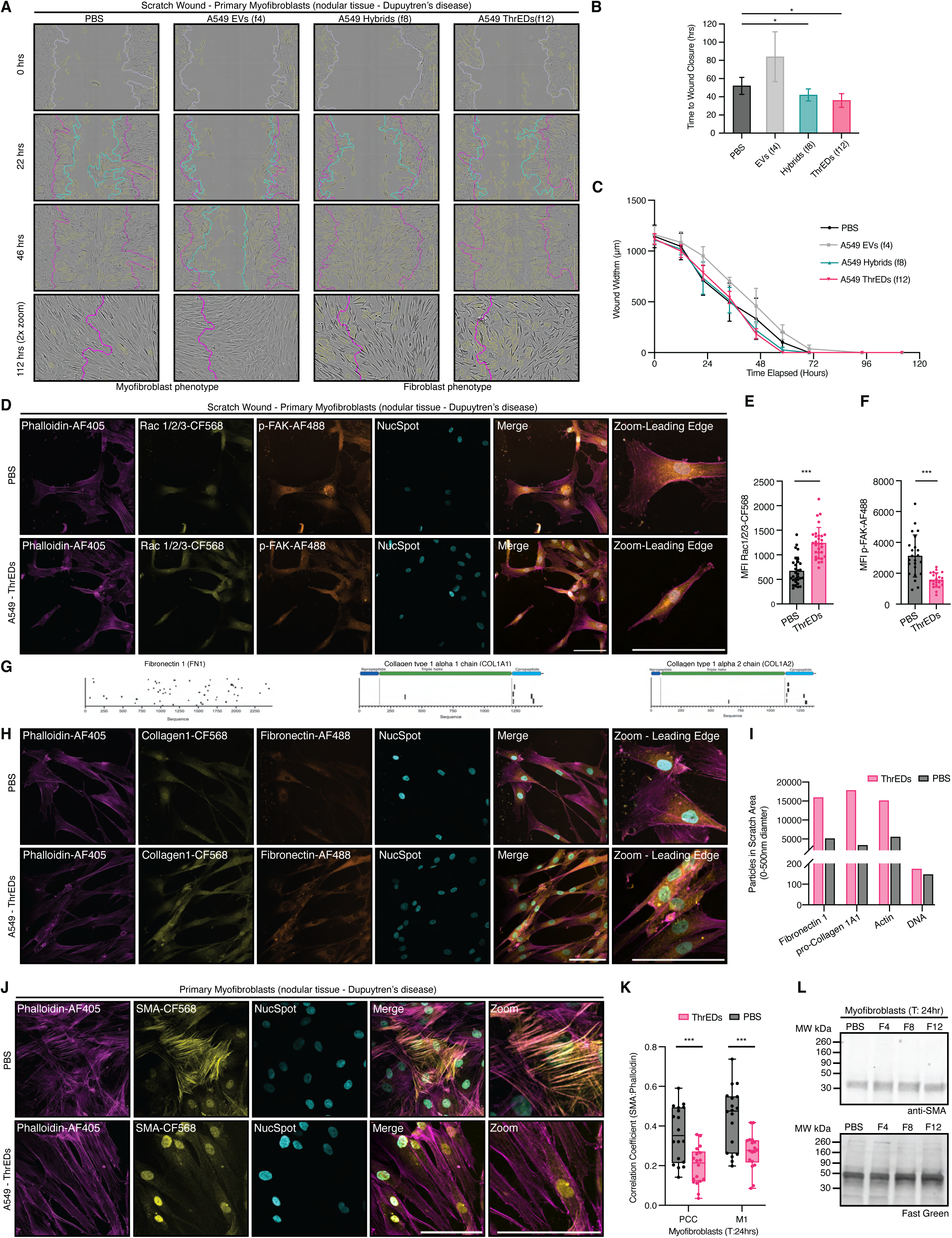
ThrEDs accelerate wound healing through cell migration and fibrotic phenotype downregulation. (A) Representative images of myofibroblast cells from Dupuytren’s disease patients treated with PBS, EVs, Hybrids and ThrEDs at T:0hr, 22 hrs, 46 hrs, and 112hrs. The initial scratch is shown with the exterior line (magenta), the scratch border is shown with the interior line (cyan), and confluence is shown throughout (yellow). The abundance of yellow borders is a proxy measure of fibrotic cell morphology. Myofibroblasts have a spread cell morphology with less “absent confluence” while fibroblasts have a spindle cell morphology with more “absent confluence”. (B) Time to wound closure in hours of A549 EPs on myofibroblasts cells. (C) Wound width for A549 EPs on myofibroblasts. (D) Confocal imaging of Rac1/2/3 and phospho-Focal Adhesion Kinase (p-FAK) 24 hrs post scratch wounding in myofibroblast cells, with associated MFI quantification in (E, F). (G) Peptide coverage on Fibronectin 1 (FN1), Collagen 1a1 (Col1a1) and Collagen 1a2 (Col1a2) proteins. (H) Collagen 1 and Fibronectin 1 confocal imaging 24 hrs post scratch wound in myofibroblast cells. (I) Quantification of the amount of FN1^+^, Col1^+^, Actin^+^ (Phalloidin), and DNA^+^ (NucSpot) particles ranging from 0-500nm in diameter deposited/present within the scratch wound gap area at 24 hrs. (J) Confocal images of myofibroblasts treated with PBS or ThrEDs strained with Phalloidin and α-smooth muscle actin (α-SMA), a fibrotic marker. (K) Associated quantification of colocalization between Phalloidin and α-SMA to determine amount of organized, fibrillar α-SMA at 24hrs post ThrEDs treatment, using Pearson (PCC) and Manders (M1). (L) Western blot probing α-SMA at 24hrs of the WCL of myofibroblasts treated with PBS, EVs, Hybrids and ThrEDs. Fast green is the loading control showing whole protein stain. Scale bars=100µm. Data represents n=3 biological replicates. n.s. P > 0.05, * P < 0.05.

Spindle cell morphology change occurs concomitantly with the loss of fibrotic α-smooth muscle actin (α-SMA) fibers in cells plated without a scratch, after 24h, persisting for ≥10 days (**Figure 6J, K**). Whole cell levels of α-SMA are not significantly different at 24h (**Figure 6L**), suggesting α-SMA fiber organization is altered. Together with decreased p-FAK, increase cellular collagen production, increased migration and cell morphology change supports a ThrED mediated downregulation of fibrotic myofibroblast phenotypes.

### ThrEDs are a novel therapeutic vehicle

A Thrombospondin-1-mClover3 chimera was used to conduct a proof-of-concept study for ThrEDs engineering. A549 release fewer EPs overall (**Figure 1E**), however a greater proportion of these EPs are ThrEDs (**Figure 2E**). As such, with a ∼35% transfection efficiency 2.8% of EPs are GFP^+^. Triton-X-100 treatment increases this proportion to 42.4% demonstrating ThrEDs incorporate the GFP fusion into the non-lipidic NVEP population (**Figure 7A)**. Iodixanol separation confirmed the presence of TSP1-GFP predominantly in peak 3 with a smaller but significant population in peak 2 (**Figure 7D**).

**Figure 7.**
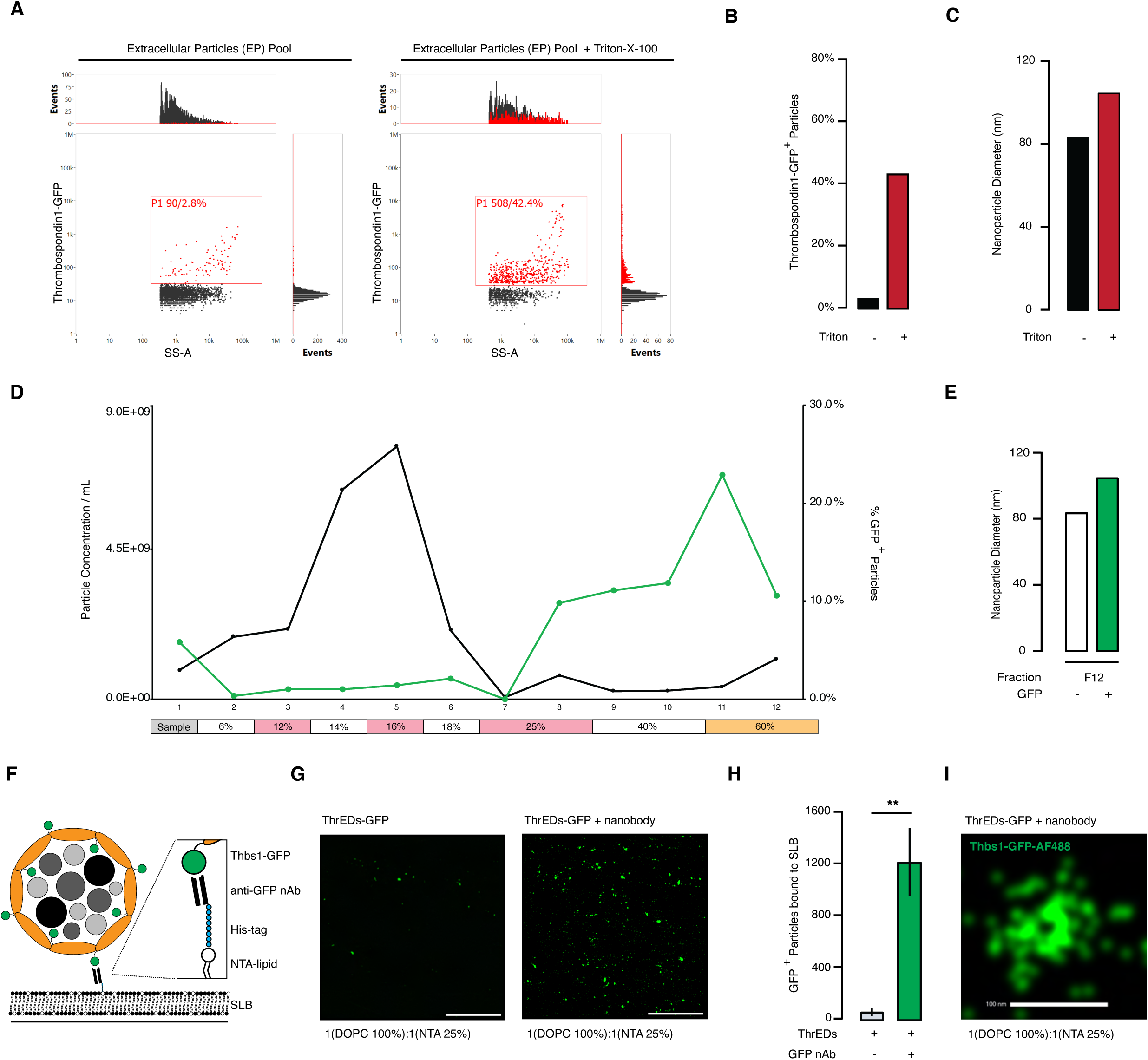
Engineering ThrEDs as a novel therapeutic vehicle. (A) EP pool from conditioned media measured using nanoflow cytometry for GFP+ particles pre and post treatment with 0.25% Triton-X-100. (B) Number of GFP+ particles and (C) their respective sizes. (D) Migration profile of EPs, isolated in 0.9mM CaCl_2_ at a 96-hour time point, following iodixanol density-based ultracentrifuge fractionation (GFP+ particle curve is the top curve for fractions 8-12). (E) Diameter of GFP+ and GFP- particles migrating to f12. (F) Diagram of anti GFP nanobody based ThrEDs capture on cell surface mimic supported lipid bilayer (SLB). (G-H) Representative confocal image micrographs of SLB capture with and without anti-GFP nanobody. Scale bar=50µm. (H) and associated quantification of particles. (I) 2D SMLM dSTORM image of a single GFP+ ThrED showing maintenance of ring structure following GFP C-terminal TSP-1 fusion. Data are presented as the mean ± SEM, n = 1 (A-E) n=3 (G-H) biological replicates, n.s. P > 0.05, * P < 0.05, ** P < 0.01, *** P < 0.001.

If the GFP at the C-terminus of TSP1 is part of the ThrEDs shell, it may be accessible to interact with an antibody attached to larger structures like cell membranes. We asked if GFP+ ThrEDs could be captured on a supported lipid bilayer (SLB) with an attached anti-GFP nanobody (**Figure 7F)**. A significant increase in GFP^+^ particle binding to the SLB in the presence of the anti-GFP nanobody was observed (**Figure 7G, H)** highlighting successful functionalization and specificity of binding despite high concentration of ThrEDs, which may have been ‘sticky’. Importantly, when SLBs were fixed and imaged by super-resolution dSTORM, ThrEDs maintain their ring structure following GFP incorporation and SLB binding highlighting ThrED stability throughout functionalization (**Figure 7I**).

## DISCUSSION

### Previously undescribed NVEP subtype in cell culture and human tissue co-isolate with EVs

EP heterogeneity has presented both a challenge and opportunity to better understand particle mediated cell-cell communication and necessitated the writing of guidelines to enhance the rigor, reproducibility and transparency of EV and EP research^41^. Despite compliant investigation of EV preparations, Thrombospondin Encapsulated Dense Particles (ThrEDs) went unnoticed. ThrEDs and EVs have similar size profiles (**Figure 1, 3, 4, S2**) and therefore co-purify by TFF, UF, common and functionalized (CaptoCore^42^) size exclusion columns. Their density likely causes co-isolation using dUC protocols.

ThrEDs may not have been previously detected as they have been dismissed as simple aggregates or protein corona^43^ due to their density^44^. Peak 2 contains both EVs and ThrEDs that may bind and function cooperatively. The hybrids have slightly stronger migration bioactivity on A549 compared to ThrEDs alone (**Figure S10**), with both mediating downregulation of the myofibroblast phenotype (**Figure 6**). ThrEDs-MMP remodeling may facilitate the EV’s ECM navigation where lipids^45^ or a transmembrane protein elicit a specific signaling cascade in the recipient cell.

Additionally, their distinct release kinetics and divalent ion stabilization requirements cause lower abundance in cell culture during EP production. EVs are typically collected after 48h, however, ThrEDs are released largely at 96h presumably as they switch to stress responsive release of proteinaceous particles over EVs (**Figure S3**). Thrombospondin accumulating at the ER, in high calcium, awaiting Calreticulin folding^46^ may spontaneously adopt a thermodynamically stable confirmation by encapsulating a condensate core of ECM related proteins as they transition through the cell’s secretory pathway^47^. The export of these dense, self-catalysing particles may be more efficient than ESCRT dependent methods^48^. ThrEDs may also be less energetically expensive on recipient cells where local calcium microenvironments induce disassembly of ThrEDs – stabilized in the ECM (high calcium) and destabilized near cells and in the cytosol (low calcium)^49^.

### ThrEDs and SMAPs function cooperatively towards a ‘repair or kill’ paradigm

The uniform protein composition regardless of source (**Figure 5**) suggests ThrEDs are a conserved particle type that increase cell survival, promoting their ubiquitous nature. Though SMAPs and ThrEDs have different functions, both contain ECM proteins, proteoglycans, glycoproteins, signaling molecules, chaperones and the unifying Thrombospondin shell^8^. As in ThrEDs, proteins in SMAPs bind to calcium for stability and function such that its absence attenuates their cytotoxicity; Perforin 1’s bioactivity is regulated by Ca^2+^ and Calreticulin^50^. Barring cytotoxicity, the GO term profiles for ThrEDs and SMAPs are similar with many cell and biological adhesion terms appearing^8^ (**Figure S6**). In specialized T cells, EPs are released with tight spatial-temporal control following APC adhesion to maximize EP potency^16^. In non-specialized cells, the stress induced release of ThrEDs facilitates the modulation of the ECM to promote healing following wounding (**Figure 6**). Generally, their dense structures protect internal, condensate cargo but are not limited by the biophysical properties of a lipid bilayer^51^ to enable cargo release (EVs). The concentrated delivery of their respective effectors leads to a rapid and robust response in recipient cells.

This class of NVEP may function cooperatively to maintain homeostasis through a ‘heal or kill’ paradigm. ThrEDs may be the first line of defense to promote tissue healing. Failure to clear the pathogenic insult would allow SMAPs to kill the perpetrator and reestablish homeostasis. For example, the formation of fibrotic tissue around a tumor due to ECM and myofibroblast deregulation restrict immune access and promote tumor growth^52^. A ThrEDs induced myofibroblast to fibroblast transition promoting migration and ECM turnover may allow immune infiltration and SMAP mediated killing of APCs. If ThrEDs cannot reestablish physiological conditions, diseased cells express moieties making them a cytotoxic target.

### Wound healing

Collagen is a critical ECM component and the most abundant protein in the body constituting ∼30% of all tissues^37^. Accordingly, the ECM is a common denominator in chronic disease as fibrosis is central to pathologies where deposition by myofibroblasts becomes excessive and generates a ‘stiff” tissue environment^53^. In ThrEDs, proteins and GO terms for both positive and negative regulation of proteolytic activity, fibril organization and cell adhesion are present (**Figure S6**). ThrEDs both accelerate cellular migration and downregulate fibrotic DD myofibroblast phenotypes (**Figure 6**).

ThrEDs proteins contain groups of functionally redundant effector protein for pro-migration and ‘soft’-matrix signaling^53^ which explains the rapid and robust responses seen with α-SMA disassembly (**Figure 6**). TGFβ, RhoA, Cdc42, ARP complex subunits 1-5 and MMPs promote migration through signaling or catalysing reorganization. Collagens I, III and XV, Fibronectin, Laminins, Lysyl oxidase provide direct architectural support and mechanotransduction signaling cues to cells. Similarly, production of ‘soft’ matrix signals reduced mechanical (fibrotic) tension in the ECM. MMPs degrade existing over deposited, stiff fibers and re-order those required for normal function. ITIH1-4 in addition to Aggrecan (**Figure S5**) promote tissue hydration through hyaluronic acid binding^54^ lowering ECM rigidity and decreasing long range transmission of mechanical stress signals^53^.

A significant increase in Fibronectin 1^+^ particles in scratch wounds suggests ThrEDs may function to nucleate the formation of new matrix against which cells can migrate for wound closure^39^. Wound healing is dependent on the re-establishment of lost ECM. Agrin^55^ and even Thrombospondin may also catalyse this formation (**Figure S5**)^56^. This suggests ThrEDs promote healing without slipping into fibrosis through an equilibrium of positive and negative regulation of a dual ThrEDs-ECM and ThrEDs-cell mechanism of action functioning both intra- and extra- cellularly.

### Biomarker potential

Of 59 proteins present in all ThrEDs, more than half have been identified in EV preparation from biological fluids (urine, plasma) as biomarkers for diseases including pancreatic and colorectal cancer^23^. Given dUC preparations, ThrEDs likely compose part of the samples. Considering ThrEDs are released as a stress response (**Figure S3**), can be isolated from plasma (**Figure 4**) and have a tight link with the ECM (**Figure 5,6**), suggests they are strong biomarker candidates.

Collagen fragments released into circulation during ECM remodeling are a strong prognostic biomarker. The Pro-C1 fragment, abundant in ThrEDs (**Figure 6**), is used to predict disease outcomes including cancer, arthritis, osteoporosis and fibrosis, including heart failure, based on proportions of formation fragments Pro-C1, Pro-C3, Pro-C4 relative to degradation fragments C1M, C3M, C4M^57^. High and low levels of fibrosis in cancer lead to drastically different prognoses due to cancer associated fibroblasts (CAFs)^58^. The early identification of a high-fibrotic phenotype is beneficial to patients. Interestingly, the Granzyme B dependent cleavage of Collagen IV (C4G fragment) was required for immune infiltration into a tumor, presenting another link between ThrEDs and SMAPs^59^. Additionally, Serglycin has high diagnostic value as a biomarker in osteosarcoma^60^.

### Engineering potential

The encapsulation of therapeutic moieties has been shown to improve their pharmacokinetic and pharmacodynamic profiles. The albumin encapsulated form of Paclitaxel (Taxol), Abraxane, is ∼130nm in diameter, has greater efficacy in some cancers due to accumulation in tumors and is better tolerated by patients^61^. The conjugation or encapsulation of therapeutics in endogenously produced protein particles present significant advantages.

We showed that Thrombospondin1-GFP form dense particles which bind to functionalized cell membrane mimics (SLBs) with a high degree of specificity, binding only to the cell membrane via the antibodies and not by sticking together or onto the membrane alone (**Figure 7**). The ability to target ThrEDs containing a densely packed core of ECM regulators to sites of injury or fibrotic disease would result in a robust but controlled response due to proteins that function on either side of an equilibrium, dampening excess deposition.

ThrEDs present significant advantages over EVs given their lack of a lipid bilayer. Fusogenic proteins are not required for the delivery of cargo to the ECM matrix, cell membrane or cytosol^51^. A change in the calcium microenvironment may be sufficient to release a high concentration of functional cargo in a localized area^49^. The potential biogenesis via condensation of protein moieties supported by charge-charge interactions would facilitate the bottom-up production of particles *ex vivo* compared to EVs. Furthermore, ThrEDs may function as more than a static therapeutic vehicle. Their abundance of ECM remodeling enzymes likely facilitates migration of particles into difficult to access areas like solid tumors. ThrEDs can therefore be used to both enhance their natural function in fibrosis and wound healing through targeting and can also be used to treat pathologies that require delivery of concentrated payloads in difficult to reach tissues.

## MATERIALS AND METHODS

### Cell culture

HEK-293T, HeLa, C2C12 and PANC1 cells were cultured in high glucose DMEM (Gibco) supplemented with 10% FBS (Gibco). Myotubes were differentiated from C2C12 by seeding cells at high density while exchanging media for DMEM supplemented with 2% Horse Serum. Cord blood Mesenchymal Stem Cells (MSC(CB)) were cultured in RPMI 1640 (Gibco) supplemented with 4ng/mL Fgf2 and 20% FBS (Gibco) and A549 cells were cultured in Ham’s F-12K (Kaighn’s) Nutrient Mixture supplemented with 15% FBS. HMEC-1 cells were cultured in MCDB131 (without L-glutamine; Gibco) supplemented with 10ng/mL EGF, 1µg/mL hydrocortisone, 10mM Glutamine and 10% FBS. HUVEC were cultured in F-12K supplemented with 0.1mg/mL heparin, 0.01mg/mL endothelial cell growth supplement (ECGS) and 10% FBS. U373 cells were cultured in EMEM supplemented with 2mM glutamine, 1% non-essential amino acids, 1mM sodium pyruvate and 10% FBS. Nalm6 cells were grown in RPMI supplemented with 10% FBS. All cell culture was performed in a humidified incubator at 37°C and 5% CO_2_.

T cells were isolated from leukapheresis reduction system (LRS) cones (‘blood cones’) obtained from healthy donors. The Oxford Radcliffe Biobank (ORB) approved access to consented and pseudonymized tissue samples under the ORB research tissue bank NHS (REC 19/SC/0173). CD8^+^ T cells were isolated by negative selection (RosetteSep Human CD8^+^ T cell enrichment cocktail, Stemcell technologies, 15023). CD8+ T cells were then either used as is (resting CD8^+^ pool) or further differentiated into CTLs using Dynabeads (Thermo, 11132D; antiCD3/anti-CD28). Resting and activated samples were grown in R10 medium (RPMI 1640, Gibco) supplemented with 1% L-Glutamine, 10% FBS, 1% Penicillin-Streptomycin, and 50 Units/mL of recombinant human IL-2 (PeproTech). For CTLs, the Dynabeads were removed after three days and cells were seeded in fresh complete R10 medium at 1E106 cells/mL for two days.

Primary fibroblast/myofibroblasts were obtained from nodular tissue from patients with Dupuytren’s disease undergoing derma fasciectomy. Informed written consent was obtained through approval of the regional ethics committee (REC 07/H0706/81), in accordance with the Declaration of Helsinki. Tissue samples were dissected into small pieces and digested in Dulbecco’s Modified Eagle Medium (DMEM) (Gibco) with type I collagenase 4mg/ml (Worthington Biochemical Corporation) + DNase I (Roche Diagnostics) for up to 3 hours at 37 °C. Digested tissue fragments were filtered through a 70-μm cell strainer, and the cell suspension was centrifuged at 1500 rpm for 10mins, washed a further 2 times in 50mls of DMEM with 5% FBS and 1% penicillin–streptomycin and the isolated cells were cultured in the same media and used until passage 5.

### Generation of constructs

Codon-optimized DNA sequences coding for human CD63, human CD81, human CD9, human TSPAN2, human TSPAN14, human PTGFRN, human PTTG1IP, human Lamp2B, human N-transferrinR1, human HSPA8, human HSP90AA1, human N-syntenin, human BASP1, human TSG101 and mClover3 were synthesized (IDT) as gene fragments and cloned downstream of the CMV promoter in a custom lentiviral backbone using PaqC1. All expression cassettes were confirmed by sequencing.

### Generation of stable cell lines

Lentiviral supernatants were generated by co-transfection of HEK-293T cells with psPAX, VSV-G and the plasmid containing each of the EV marker proteins fused to mClover3 via a flexible linker using 1mg/mL PEI. 48 hours post transfection lentiviral supernatants were collected, cleared at 500 x g at 4°C and filter sterilized using a 0.45µm filter. Supernatants were aliquoted and frozen at −80°C.

For the generation of stable cell lines, HEK-293T, HeLa, A549 and MSC (CB) cells were plated in 6 well plates (Corning). Twenty-four hours later, lentiviral supernatant was added with 10µg/mL polybrene. Twenty-four hours later lentiviral supernatant was removed, cells were allowed to recover for forty-eight hours prior to selection with 2µg/mL puromycin. Cells were passaged at least five times under puromycin selection.

### Transient transfection

HEK-293T cells were plated at 60% confluence into a 12-well plate (Corning) or a 15-cm dish (Corning) for SEC analyses. The next day, 1.4µg total DNA (12-well) and 50µg total DNA (15cm plate) was combined with OptiMEM. Then 3.5µg (12-well) and 125µg (15cm plate) of PEI was combined with OptiMEM in a separate tube. Opti-DNA and Opti-PEI solutions were combined in a 1:1 ratio, vortexed, and allowed to sit at RT for 20 minutes. The mixture was added to each plate in a dropwise manner. The transfection media was changed to OptiMEM the following day, and EPs were harvested 48 hrs after medium change when cells had reached 90% confluence.

### EP isolation

CM protocol – Stable or wild type cells were plated into a 12-well plate at 70-80% confluence. The next day the media was exchanged for serum depleted OptiMEM. 48 hours later, Conditioned media (CM) was collected from 12-well plates. First, CM was pre-cleared at 500xg for 10 mins at 4°C. The supernatant was then subjected to 4000xg for 20 mins at 4°C. CM EPs were then used for downstream nanoflow cytometry analyses.

SEC protocol – Conditioned media (CM), generated as above, was collected from 15cm plates. First, CM was pre-cleared at 500xg for 10 mins at 4°C. The supernatant was then subjected to 4000xg for 20 mins at 4°C. The resulting supernatant was concentrated using tangential flow filtration (VivaFlow 100 kDa MWCO) down to around 10mL. Concentrated media was then subjected to spin filtration using Millipore Amico 100 kDa MWCO to further concentrate media to allow for injection on size exclusion column Sepharose 4 Fast Flow resin in a Tricorn 10/300 column. EPs were eluted in low protein binding tubes (Corning, Pyrex 12X75mm disposable rimless culture tubes) and combined for downstream analyses.

### ThrED particle isolations

EPs were isolated as above using the SEC protocol then loaded onto an iodixanol gradient as previously described^33^. All iodixanol solutions were produced using 60% stock OptiPrep (Sigma-Aldrich), filter sterilized (0.22µm) MilliQ H_2_0, Phenol Red (0.5% Sigma-Aldrich), filter sterilized (0.22µm) 5xPBS-MK (250 ml PBS 10X, 2.5 ml 1M MgCl^2^, 6.25 ml 1M KCl, H_2_O up to 500 ml), and 5M NaCl for the 6% solution only. 6%, 12%, 14%, 16%, 18%, 25%, 40% and 60% iodixanol solutions were prepared. Then poured as follows, 2mL 60%, 2mL 40%, 2mL 25%, 1mL 18%, 1mL 16%, 1mL 14%, 1mL 12% and 1mL 6%. 0.5mL of concentrated sample volume is added carefully to the top of the Beckmann-Coulter 13.2mL thinwall ultra-clear ultracentrifuge tube. Samples were spun overnight for 16 hrs at 28,500 RPM and 4°C in a SW41Ti rotor. The next day, fractions were collected from the top of the gradient. Each fraction was spun separately and iodixanol was exchanged for the appropriate buffer using a Millipore Amicon 100 kDa MWCO spin filter.

### ThrEDs particle isolation from plasma

Peripheral blood from healthy, fasted donors was acquired under the ethics license REC 11/H0711/7 (University of Oxford). To obtain plasma, blood was collected in BD Vacutainer® ACD A Blood Collection Tubes (Trisodium Citrate, 22.0 g/L, Citric Acid, 8.0 g/L, and Dextrose, 24.5 g/L). Blood was transferred to centrifuge tubes and spun at 2,500xg for 15min at RT. The upper plasma layer was transferred to fresh tubes and spun again at 2,500xg for 15min and RT with no break. The supernatant was transferred to a free tube and spun again to generate platelet free plasma “PFP”. PFP was then diluted in 1XPBS with 25mM HEPES (PBS-H) 1:10 to be used within 2 weeks or frozen undiluted at −80°C for up to 2 years.

Diluted PFP was then loaded into open top Thinwall Ultra-Clear tubes 14x89mm 13.2mL (Beckman Coulter; cat. 344059) and loaded into a SW 41 Ti swinging bucket rotor and spun for 4 hours at 150,000xg, 4°C. The supernatant is then carefully aspirated. The pellet is diluted again in 1XPBS-H and spun with the same settings a second time. The resulting pellet is then run on a size exclusion column.

The concentrated sample is then depleted of albumin using the Abcam albumin depletion kit (Abcam; ab241023). Albumin depleted EPs were then loaded onto an iodixanol gradient as above.

### Nanoflow cytometry

EP samples from crude CM, SEC and iodixanol purifications were analyzed by the nanoFCM. The instrument was calibrated using QC beads (200nm PE and AF488 fluorophore conjugated polystyrene beads) and Sizing beads (S16M-exo beads from 40-200nm in diameter silica nanosphere cocktail). EP samples were diluted to achieve the optimal 2,000-12,000 particle counts/min.

1E8-1E10 particles were used for all staining with antibodies listed in Table 1. Samples were incubated overnight at 4°C and if necessary, a secondary antibody was added and incubated for 2-4 hrs. Samples were diluted 1:100 prior to measurement on nanoFCM to remove background signal or purified using SEC.

**Table 1.** Antibodies and Stains used for WB, FC, nFC and IF (confocal, dSTORM)

| Protein Name | Antibody/Stain | Dilution | Application |
| --- | --- | --- | --- |
| CD63 | Rabbit anti-CD63-PE | 1:100 | Nanoflow cytometry |
| ApoE | Rabbit anti-APOE BioRad | 1:100 | Nanoflow cytometry |
| Thbs1 | Mouse anti-Thrombospondin 1 Santa Cruz - CF568 | 1:100 | Nanoflow cytometry, dSTORM |
| Thbs1 | Mouse anti-Thrombospondin 1 Santa Cruz – Atto 488 | 1:100 | dSTORM |
| Thbs1 | Mouse anti-Thrombospondin 1 Santa Cruz | 1:1000 | Western Blot |
| Srgn | Rabbit anti-Serglycin Santa Cruz | 1:100, 1:1000 | Nanoflow cytometry, Western Blot |
| Srgn | Rabbit anti-Serglycin – AF647 Santa Cruz | 1:100 | Nanoflow cytometry, dSTORM |
| Thbs5 | Rabbit anti-Thrombospondin 5 (COMP) | 1:100 | Nanoflow cytometry, dSTORM |
| GrzmB | Rabbit anti-Granzyme B AF647 | 1:100 | Nanoflow cytometry |
| GrzmB | Rabbit anti-Granzyme B Cell Signalling Technologies | 1:1000 | Western Blot |
| Prfn1 | Mouse anti-Perforin 1 AF488 | 1:100 | Nanoflow cytometry |
| Prfn1 | Mouse anti-Perforin 1 Santa Cruz | 1:1000 | Western Blot |
| Casp3 | Rabbit anti-Cleaved Caspase 3 Cell Signaling Technologies | 1:500 | Flow cytometry |
| Actin | Phalloidin AF405 plus | 1:40 | Confocal |
| SMA | Rabbit anti-α smooth muscle actin Abcam | 1:2000 | Western blot |
| SMA | Rabbit anti-Smooth Muscle Actin Abcam – CF568 | 1:40 | Confocal |
| Rac1/2/3 | Rabbit anti-Rac1/2/3 brand – CF568 | 1:40 | Confocal |
| p-FAK | Rabbit anti-phospo-Focal Adhesion Kinase Thermo – AF488 | 1:40 | Confocal |
| Collagen 1A1 | Mouse anti-pro Collagen1A1 Thermo – CF568 | 1:40 | Confocal |
| Fibronectin 1 | Mouse anti-Fibronectin 1 – Dylight488 | 1:40 | Confocal |
| GFP | Mouse anti-GFP Biolegend | 1:100 | dSTORM |
| - | Secondary 488 | 1:10,000 | Nanoflow cytometry, dSTORM |
| - | Secondary 647 | 1:10,000 | Nanoflow cytometry, dSTORM |
| - | Goat anti-Rabbit Alexa Fluor 647 | 1:10,000 | Nanoflow cytometry, dSTORM |
| - | Secondary rabbit HRP | 1:10,000 | Western Blot |
| - | Secondary mouse HRP | 1:10,000 | Western Blot |
| - | Goat anti- rabbit LiCOR 680-700 | 1:25,000 | Western Blot |
| Lipid Membrane | Cell Mask Green | 1:200 | Nanoflow Cytometry |
| Lipid Membrane | Cell Mask Far Red | 1:200 | Nanoflow Cytometry |
| Lipid Membrane | DiR | 1:200,1:100,1:50 | dSTORM |
| Glc-Nac Protein | Wheat Germ Agglutinin – AF488 | 1:200 | Nanoflow cytometry, dSTORM |
| Glc-Nac Protein | Wheat Germ Agglutinin – AF568 | 1:200 | dSTORM |
| Glc-Nac Protein | Wheat Germ Agglutinin – CF568 | 1:200 | dSTORM |
| Glc-Nac Protein | Wheat Germ Agglutinin – AF647 | 1:200 | Nanoflow cytometry |
| Live/dead | Live-dead stain Aqua blue | 1:400 | Flow cytometry |

EDTA treatment of individual iodixanol fractions was carried out by adding EDTA to a final concentration of 0.5mM and allowed to rotate overnight at 4°C. Samples were run as usual the next day. For EP detergent treatments, Triton-X-100 was added to a final concentration of 0.25%, vortexed for 30 sec then re-run immediately on the nanoFCM to determine the proportion of remaining, detergent resistant particles.

### Western blotting

EP fractions were pooled and sodium deoxycholate (ThermoFisher) was added to a final concentration of 1%, samples were rotated for 15 minutes. NuPage LDS Sample Buffer (4x; Thermo Fischer) supplemented with NuPage Sample Reducing Agent (10x; Thermo Fisher) was added to a final 1x concentration then boiled at 95°C for 5 minutes. ThrEDs were prepared similarly, but without the initial sodium deoxycholate detergent.

Whole cell lysate (WCL) preparations were generated by washing a 15cm plate in 1xPBS then adding 1xRIPA buffer supplemented with protease inhibitors (Merck, cOmplete, Mini, EDTA-free Protease inhibitor cocktail). Samples were aliquoted and sonicated for 2 minutes. LDS sample buffer and reducing agent were added as above then samples were boiled at 95°C for 5 minutes. Protein concentrations for western blotting were normalized using a BCA assay. 6µg of EP preparations and WCL were added to each well (∼1E9 particles) for cell line western blots. Western blots using human tissue, including DD patients myofibroblasts and donor matched CD8+ rest and CTLs samples were precipitated using 10% TCA for 1 hour then washed with 0.1% TCA prior to resuspension in LDS buffer. 10µg of protein were loaded per well for human tissue.

Samples were run on a Bis-Tris 4-12% SDS PAGE gel using MOPS buffer. Proteins were then transferred onto a PVDF membrane activated in 100% methanol. The gel was transferred using NuPAGE transfer buffer with 20% methanol. Membranes were blocked for 1 hours in 5% skim milk PBS-T at RT then incubated overnight at 4°C with relevant antibody in 5% skim milk PBS-T. Membranes were then washed in PBS-T prior to incubation in secondary antibody, for 1 hour at RT. Membranes were washed again in PBS-T prior to visualization. ECL (BioRad Clarity Western ECL) was used for HRP secondary antibodies and imaged on the ChemiDoc using chemiluminescence. LiCOR imaging system was used to visualize LICOR secondary antibodies.

### Confocal microscopy

Confocal images were acquired with the Zeiss 980 Airyscan 2 microscope using 20x (Plan-Apochromat 20x/0.8, air), 40x (EC Plan-Neofluar 40x/1.30 Oil DIC) and 63x (Plan-Apochromat 63x/1.4 Oil DIC) objectives. 405nm, 445nm, 488nm, 514nm and 649nm diode lasers and 561nm and 594nm DPSS lasers were used to image samples. Spectral detector 32 channels GaAsP PMT plus 2 channels MA-PM and Axiocam 506 mono detectors were used. Images were taken with 4x averaging to achieve optimal signal to noise ratio, exposure time and laser power settings were kept consistent between all samples to ensure accurate representations and quantification.

For assays with immunofluorescence staining, A549 cells and primary myofibroblasts were seeded at a confluence of ∼90-95% into µ-Slide 8 Well high imaging chamber with a #1.5 Polymer Coverslip (IBIDI cat. 80806). Cells seeded the previous day were administered a scratch wound 24 hours later in A549 and 48 hours later in myofibroblasts to allow complete adherence. 24 hours following the scratch cells were washed with 1XPBS twice and fixed with 4% PFA.

Immunofluorescence samples were all stained as follows. Cells were blocked for 1 hour in Blocking Buffer (1X PBS, 5% normal goat serum, 0.3% Triton-X-100) at RT, then washed 3 times with 1XPBS. All primary antibodies were pre-conjugated to fluorophores. Antibodies were diluted in Antibody Dilution Buffer (1XPBS, 5% BSA, 0.3% Triton-X-100) as follows; SMA-CF568 (1:50), Rac1/2/3-CF568 (1:40), p-FAK-AF488 (1:40), proCol1A1-CF568 (1:40), Fibronectin-AF488 (1:40). Samples were stained overnight at 4°C then washed 3x with 1XPBS prior to staining with Phalloidin-AF405 Plus (1:40) and NucSpot 650/665 (1:1000) in sequence for 1 hour each with a 1XPBS wash in between.

### dSTORM microscopy

The Oxford NanoImager (ONI) microscope was used to generate 2D SMLM (single molecule localization microscopy) images using dSTORM (direct Stochastic Optical Reconstruction Microscopy). 1E8-1E10 particles were diluted to a final volume of 300µL (1xPBS for EVs or Ca^2+^ buffer for protein particles). 1-4µL of 2.5mg/mL DiR and 1µL of 2mg/mL WGA-AF568/647 or 2µL of 1mg/mL WGA-AF488 were incubated overnight while rotating at 4°C. Alternatively, 4µL of 2.5mg/mL DiR, 2µL of mouse anti-human Serglycin (C-11; Santa Cruz Biotechnology, sc-374657) and 1µL of TSP1-CF568 (Thermo Fisher, clone A6.1, MA513398) were used together. 1µL of goat anti-mouse 488 secondary antibody was added after primary overnight incubation and left for 2 hrs. After staining, samples were re-purified by SEC to reduce noise in the imaging chamber and to remove all soluble proteins present, avoiding false positive detection of protein particles due to protein aggregate staining with WGA, Thrombospondin-1 or Serglycin.

Imaging chamber was prepared the same day as imaging using Ibidi VI 0.4 channel slide (#80608) and 1.5H 25x75mm cover slips. Poly-D-lysine was added and incubated for 2 hrs. Each chamber was washed with 1xPBS 3 times. Stained and re-purified samples (∼1E8-1E9 particles) were added to each chamber. Samples were allowed to settle overnight at 4°C (at least 4 hours). Unbound particles were washed out with 1xPBS 3 times. Samples were imaged in dSTORM buffer (320µL 1xPBS, 40µL 50% glucose, 40µL 2-Mercaptoethylamine•HCl (2-MEA), 4µL glucose-oxidase). First, 640nm laser was used to excite DiR or AF647, then the 561nm laser was used to excite the Alexa568 dye or CF568, followed by the 488 laser used to excite the Alexa488 dye or mClover3 bound to AF488 anti GFP FAb. 3,000-10,000 frames were acquired per fluorophore.

### Proteomics

#### Sample preparation

ThrED and whole cell lysate samples were prepared for mass spectrometry as above. Sodium Dodecyl Sulfate was added to samples to a final concentration of 5% before they were reduced and alkylated using TCEP (10mM) and Iodoacetamide (50mM) respectively. Phosphoric acid was added to a final concentration of 2.5% followed by 7 volumes of S-trap binding buffer (90% methanol in 100mM TEAB). Samples were then loaded onto s-trap columns (ProtiFi), following washes with S-trap binding buffer, trypsin in 50mM TEAB was added (ratio 1:30) and samples were digested overnight. Samples were eluted with 50mM TEAB, followed by 0.2% formic acid and then finally with 0.2% formic acid in 50% acetonitrile. Elutions were pooled and dried down and resuspended in 0.1% formic acid.

#### Mass Spectrometry – LC-MS/MS Analysis

Liquid chromatography–tandem mass spectrometry (LC–MS/MS) was used to analyse the tryptic peptides using the Vanquish Neo UHPLC connected to the Orbitrap Ascend Mass Spectrometer (all Thermo Fisher Scientific) with an EASY-Spray Source. The Vanquish Neo was operated in “Trap and Elute” mode using a PepMap Neo trap (5 μm, 300 μm x 5 mm; Thermo Fisher) and separated using an EASY-SPRAY PepMapNeo column (50 cm x 75 um, 1500 bar; Thermo Fisher) heated at 50C. Mobile phase A and B were 0.1% Formic acid in water (LC-MS Optima grade) and 0.1% Formic acid in Acetonitrile (LC-MS Optima grade), respectively. Tryptic peptides were trapped and separated over a 75 min gradient, going from 2 to 18% B in 40 min, to 35% in 20 min, up to 99% B in 1 min and then staying at 99% for 14 min. The flow rate was maintained at 300 nL/min throughout the gradient. Data acquisition was carried out in data-independent mode (DIA) similar to previous studies^62–64^. MS1 scans were collected in the orbitrap at a resolving power of 45K at m/z 200 over m/z range of 350 to 1,650 m/z. The MS1 normalized automatic gain control was set at 125% (5e5ions) with a maximum injection time of 91ms and a radio frequency lens at 30%. DIA MS2 scans were then acquired using the tMSn scan function at 30K orbitrap resolution over 40 scan windows with variable width, with a normalized automatic gain control target of 1,000%, maximum injection time set to auto and a 30% collision energy.

#### Data analysis

Raw data was searched using DIA-NN (v1.8.1) against UniProt/SwissProt human database (2024) with added decoys and known contaminants which contained a total of 40936 sequences (50% decoys). Searches used a maximum false discovery rate of 1%. Carbamidomethyl was added as a fixed modification to cysteine residues (+57.021Da). A fully tryptic search was conducted with 2 missed cleavages allowed. Data was exported and onward analysis was conducted in Perseus (v2.0.11) and R (v4.3.2). GO pathway analysis was conducted using Enrichr with a background of all proteins detected in this study^65^. Revigo (v1.8.1) was used to remove redundant GO terms^66^. For polarity and ECM related GO terms descriptions were filtered for specific key words relating to these topics. STRING (v12) was used for network analysis^67^. Matrisome analysis was done using MatrisomeDB as a reference database^68^.

### Cytotoxicity Assay

Cytotoxicity of proteinaceous particles (ThrEDs and SMAPs) was assessed by measuring the viability of Nalm6 cells. 2.0E4 cells were combined at a 1:100 ratio with particles in a round bottom 96-well plate for 72 hours before harvesting. Cells were stained with Live/Dead^TM^ Fixable eF780 dye (Invitrogen) followed by 4% PFA fixation. After fixation, cells were permeabilised with 0.1% Saponin w/v HBSS for 20 mins at RT. Cells were then blocked with 5% BSA w/v HBSS for 1 hour at RT prior to staining with 1 µg/ml anti-cleaved caspase-3 for 1 hour at 4 °C. The cells were then washed twice with 200 µl 0.02% Saponin w/v HBSS and analysed on a BD Fortessa. Data was analysed using FlowJo v10.10.0.

### Supported Lipid Bilayer (SLB) Targeting

Glass coverslips (Nexterion) were plasma cleaned and mounted onto µ-Slide VI 0.4 (six channel chambers). DOPC (1,2-dioleoyl-sn-glycero-3-phosphocholine) and NTA (1,2-di-(9Z-octadecenoyl)-sn-glycero-3-[(N- (5-amino-1-carboxypentyl) iminodiacetic acid) succinyl] (nickel salt)) lipids were combined and added to each chamber. NTA lipids were used at a final concentration of 12.5% to bind His tagged proteins. Channels are washed with HBS+0.1% BSA in between each of the following steps. Following removal of free lipids, the channels were blocked with HBS+ 3% BSA for 30 minutes. Then 8xHis-tagged GFP-nAb are added to the channel and left to equilibrate, binding with NTA lipids, for 1 hour. After removing unbound nAb, Thbs1-GFP ThrEDs were added for 1 hour. Unbound ThrEDs were washed away thoroughly.

IBIDI channels were first imaged by confocal microscopy to determine degree of Thbs1-GFP – GFP-nAb binding. Then, samples were fixed with 4% PFA and 0.25% Gluteraldehyde for 1 hour at 4°C. After washing fixation medium out of the channels, the samples were imaged by dSTORM to determine the structure of particles bound to the SLB.

### Electron Microscopy

ThrEDs were prepared for negative stain electron microscopy using the methods described above. Final suspension buffers used for EM were PBS, MilliQ + 4.5mM CaCl_2_, MilliQ + 0.5mM EDTA.

Freshly glow-discharged TEM grids (C267, TAAB) were placed carbon-side down onto 10 µl droplets of sample and incubated for 2 minutes. The grids were then blotted and incubated for 10 seconds on a 20 µl droplet of aqueous uranyl acetate 2% (w/v), then blotted and allowed to air dry. Grids were imaged using a JEOL Flash 120kV TEM equipped with a Gatan Rio camera.

### Wounding and migration assay

The evening before the wounding assay, 34.0E4 to 38.0E4 A549, primary fibroblasts recipient cells were plated into a Incucyte Imagelock 96-well microplate (BA-04856). The next day, when cells were 100% confluent, the Incucyte 96-Well Woundmaker tool was used to generate 96 uniform scratches. After washing 3 times with PBS, EPs were added. The plate was then added into the Incucyte and imaged using the 20x objective every 2 hours over 120 hours to capture wound closure. Wound healing was then assessed using the Incucyte Scratch Wound Analysis Software Module.

### Fibrotic Phenotype Assay

The ability of ThrEDs to downregulate fibrotic marker expression and phenotype in cells was conducted by plating primary myofibroblasts at ∼90-95% into µ-Slide 8 Well high imaging chamber with a #1.5 Polymer Coverslip (IBIDI cat. 80806). 48 hours later, 1E8 extracellular particles were added into each well and left for 24 hours for IF imaging or up to 10 days to observe long term phenotype. Whole cell lysate was also collected following 24 hours using 1XRIPA with protease inhibitors.

### Image analysis

The CODI platform was used to analyse dSTORM images acquired with the ONI system, as described previously^10^.

Image J was used to analyse confocal images acquired using the Zeiss 980 Confocal and western blot images. Incucyte images were analysed using integrated Incucyte Scratch Wound software.

### Statistical analysis

Excel and Prism (v.10.4.0) was used to perform statistical analysis and generate graphs. Differences between two groups were assessed by Student’s t-test.

## Supplemental Figure Legends

**Figure S1.**
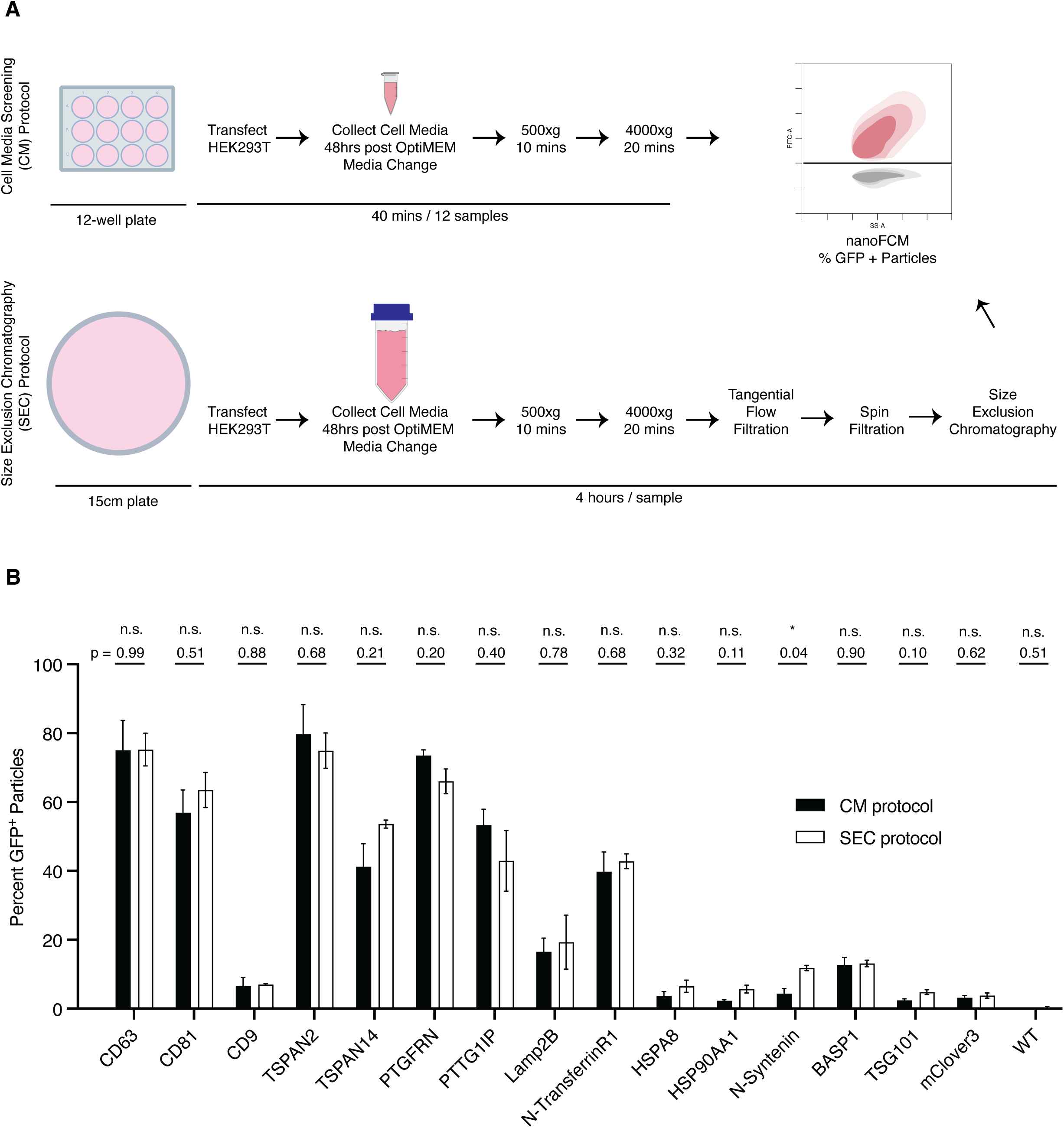
Validating an EV marker screening assay for single particle analysis. (A) Schematic representing the EV marker screening assay protocol. (B) Percentage of GFP+ (mClover3) EVs from transiently transfected HEK-293T cells isolated by CM or SEC protocols. Data are presented as the mean ± SEM, n = 3 biological replicates, n.s. P > 0.05, * P < 0.05.

**Figure S2.**
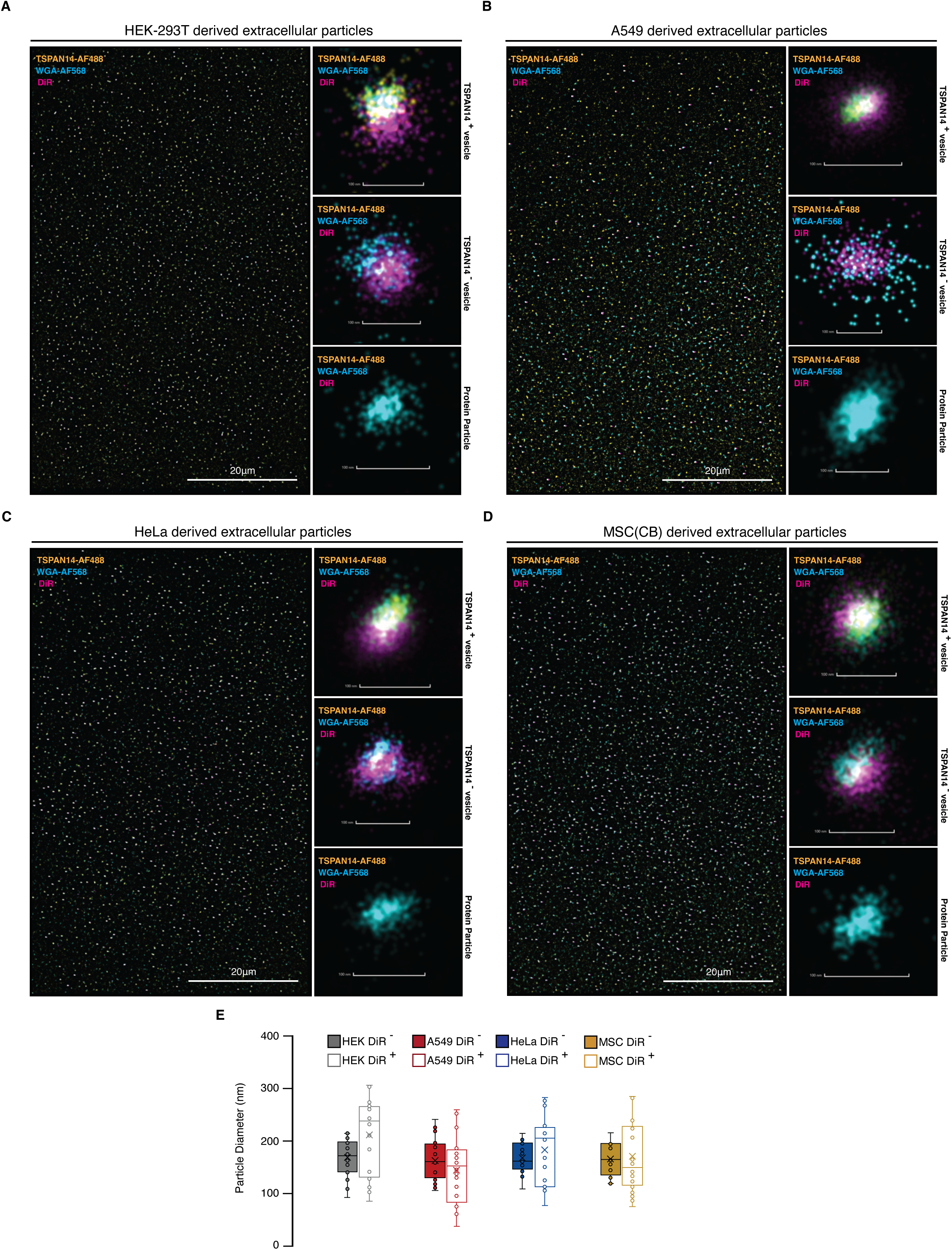
EP subpopulation dSTORM imaging confirms presence of glycosylated NVEPs. Single vesicle 2D SMLM dSTORM image analysis of nanoparticles from TSPAN14-mClover3 expressing stable cell lines in (A) HEK-293T, (B) A549, (C) HeLa, and (D) MSC(CB). Nanoparticles isolated by SEC are stained with DiR (far red membrane intercalating dye), WGA-AF568 (wheat germ agglutinin; binds N-linked glycosylated protein residues) to detect glycosylation PTMs, and anti-GFP-AF488. Scale bar 20μm in full field of view and 100nm in single particle zoom panels. Representative images of TSPAN14+ EVs, TSPAN14– EVs, and WGA+ spherical protein particles are shown. (E) Box plot of particle diameter (nm) calculated using the ONI CODI software are shown for EVs (DiR+ particles) and protein particles (DiR– particles) in each cell line (pair of bars from left to right correspond to HEK, A549, HeLa and MSC). Data represents n=20 nanoparticles per group.

**Figure S3.**
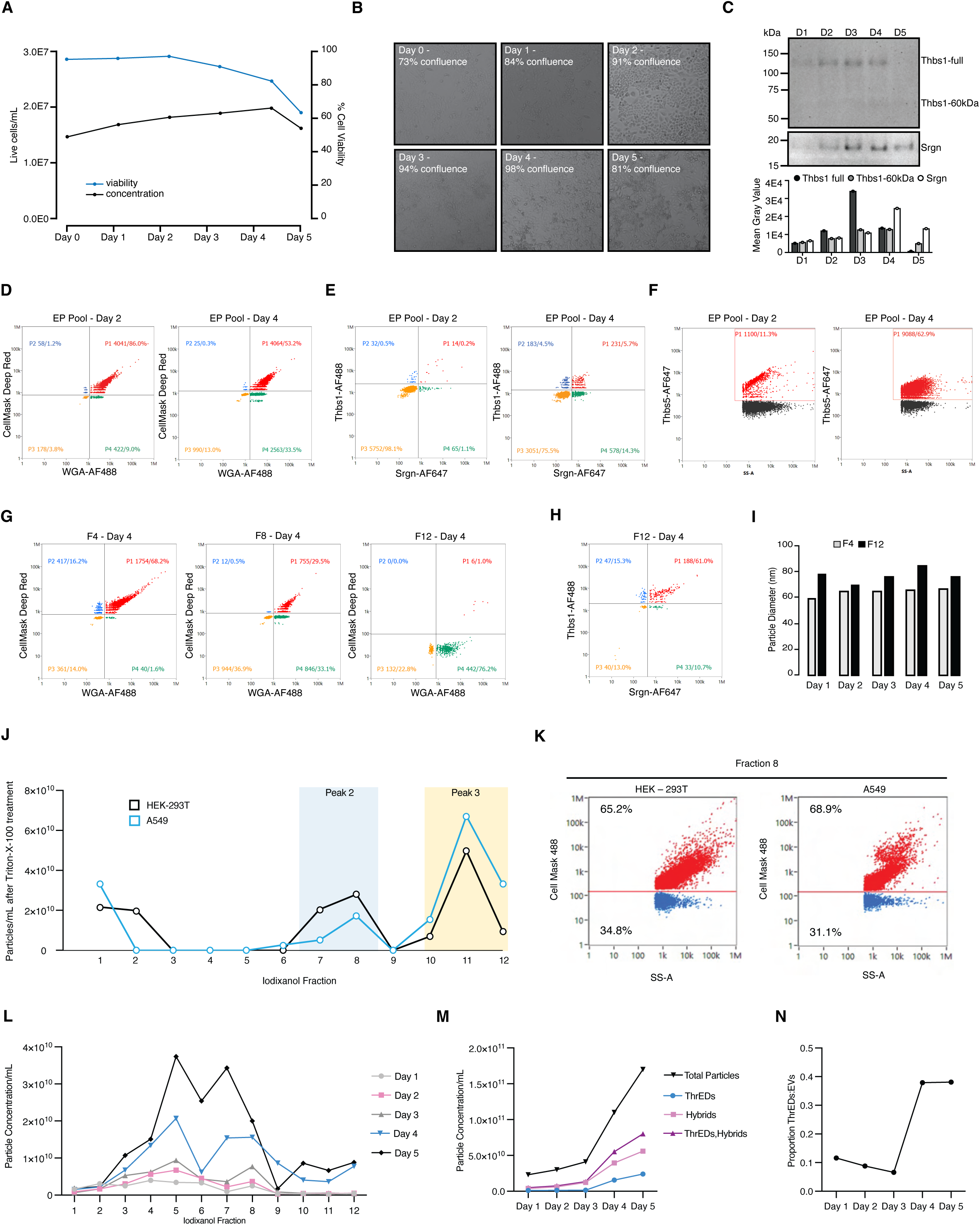
Release kinetics of Thrombospondin Encapsulated Dense particles. (A) Plot of A549 live cell concentration in cells/mL (bottom curve) and viability (top curve) at timepoints across 5 days of collection with (B) representative micrographs of cell confluence and morphology at each day. (C) Western blot whole cell lysates from day 1 through 5 probing Thrombospondin 1 and Serglycin with associated quantification of the mean grey value of each band. (D) Nanoflow cytometry analysis of extracellular particles pools from conditioned media at days 2 and 4 probing protein (WGA) and lipid (Cell Mask) biomolecules (E) TSP1 and SRGN and (F) COMP. (G) Nanoflow cytometry analysis of f4, F8 and f12 probing protein (WGA) and lipid (Cell Mask) biomolecules and (H) f12 probing Thbs1 and Srgn, all at the day 4 time point. (I) Size profiles of EVs in f4 and ThrEDs in f12 determined by side scatter using nanoflow cytometry across time points day 1 to 5. (J) EPs were isolated in 4.5mM calcium and then treated with 1% Triton-X-100. The EP Migration profile was achieved following iodixanol density-based ultracentrifuge fractionation for Triton-X-100 treated samples. (K) Following a typical protocol without Triton-X-100, f8 fractions from HEK-293T and A549 were stained for Cell Mask which labelled lipids (EVs), showing two populations in f8. (L) Migration profile of Extracellular Particles (EP), isolated in isosmotic buffer with 4.5mM CaCl_2_ at time points day 1 through 5, following iodixanol density-based ultracentrifuge fractionation. (M) Concentrations of f1-12 total particles, f12+11 ThrEDs, f7+8 Hybrids, across time points and (N) the proportion of f11+12 ThrEDs relative to the proportion of f4+5 EVs across time points.

**Figure S4.**
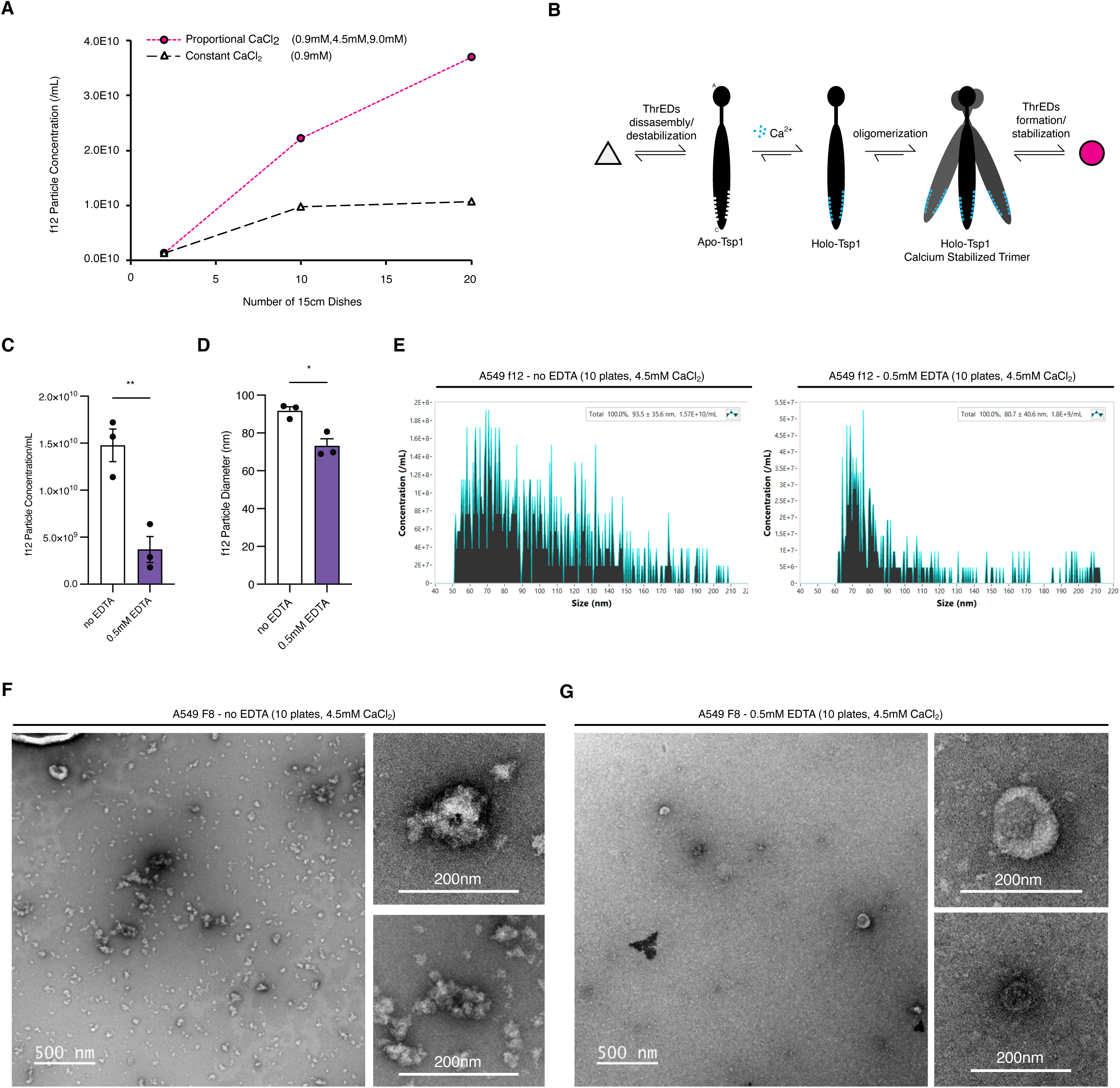
Calcium is critical in the formation and stabilization of ThrEDs. (A) The effect of increasing amount of CaCl_2_ in an isosmotic buffer on ThrEDs f12 particle concentration. Samples were isolated from 2 plates, 10 plates and 20 plates, purified with constant calcium (0.9mM CaCl_2_) or proportional amounts of calcium at 0.9mM, 4.5mM and 9mM for 2, 10 and 20 plates, respectively. (B) Diagram representing the homo-trimerization of Thrombospondin 1 induced by calcium and the effect on ThrEDs stability equilibrating between low concentration and destabilized and high concertation and stable. (C) f12 particle concentration following treatment with 0.5mM EDTA and the (D) associated change in particle size with raw size distribution plots shown in (E). (C-E) were generated from 10 plates isolated in 4.5mM CaCl2. (F-G) Demonstrate the effect of calcium stabilization on ThrEDs morphology in f8 before and after EDTA treatment. Calcium stabilization of Thrombospondin subunits is seen in (F) with dense particles abundant among EVs (hybrid population) and the loss of large ThrEDs following EDTA treatment while EVs morphology and concentration remain unchanged (G). n.s. P > 0.05, * P < 0.05, ** P < 0.01.(A) n=1, (C-G) n=3 biological replicates.

**Figure S5.**
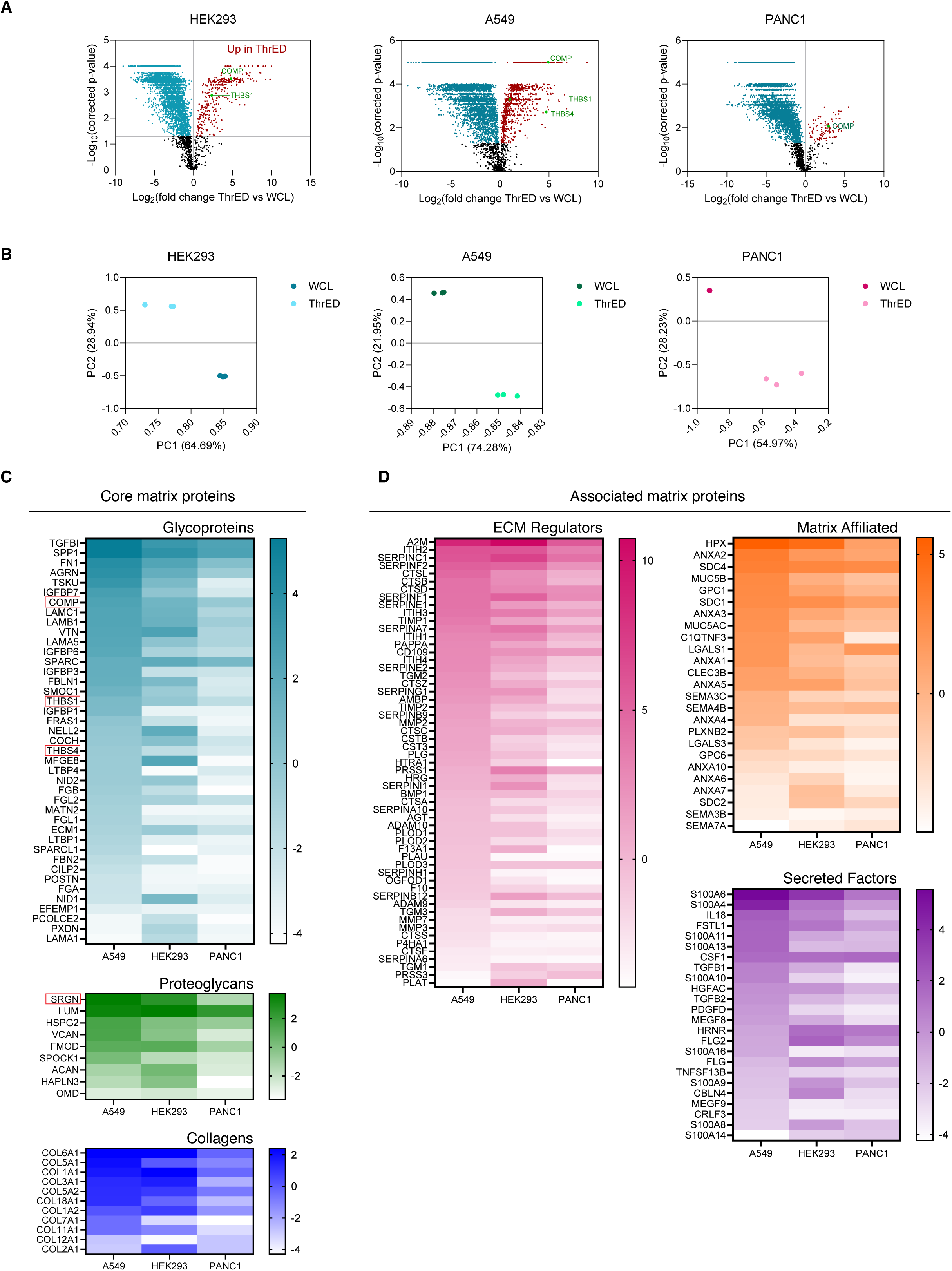
ThrED proteomics matrix highlight core and associated matrix proteins. (A) Volcano plots of proteins identified in ThrEDs. Red and blue labelled proteins indicate those significantly upregulated in ThrEDs or whole cell lysate, respectively. (B) PCA plots for the proteins identified in f11+12 ThrEDs and whole cell lysates of HEK293, A549, and PANC1 cells. (C-D) Heat maps of proteins quantified (Log_2_ abundance) in ThrEDs in A549, HEK293, PANC1 sorted into protein classes and functional groupings as determined by MatrisomeDB. n=3 biological replicates.

**Figure S6.**
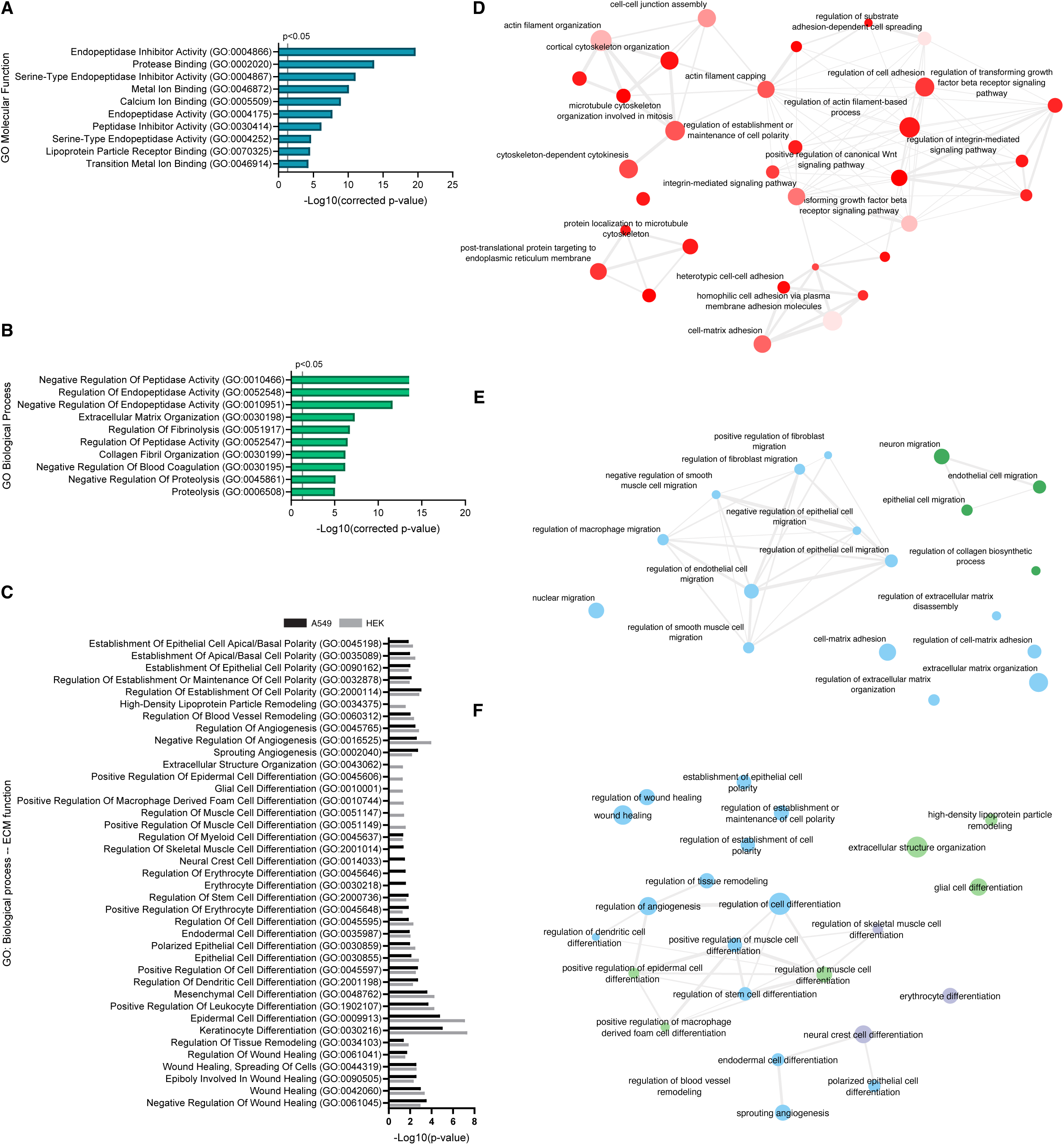
GO term analysis of ThrED proteins have complementary and opposing biological and molecular ECM function. Top ten GO terms from proteins identified in at least 2 ThrEDs sources of HEK, A549, PANC1 for (A) Molecular Function, (B) Biological Process, (C) Significant GO terms related to the formation of the ECM from proteins identified in ThrED particles in HEK and A549. (D) GO term network analysis of significant, non-redundant polarity terms identified in A549 whole cell lysates. The nodes are coloured by p-value where darker red indicated increased confidence, the size of each dot represents the number of annotations of the GO term. (E) GO term network analysis on ThrED proteins from HEK and A549 involved in general ECM biology and ECM formation and (F) ECM function. Blue points ae present in both HEK and A549. Green points are only in HEK and purple points are only in A549. The size of each dot describes the number of annotations for that GO term. n=3 biological replicates.

**Figure S7.**
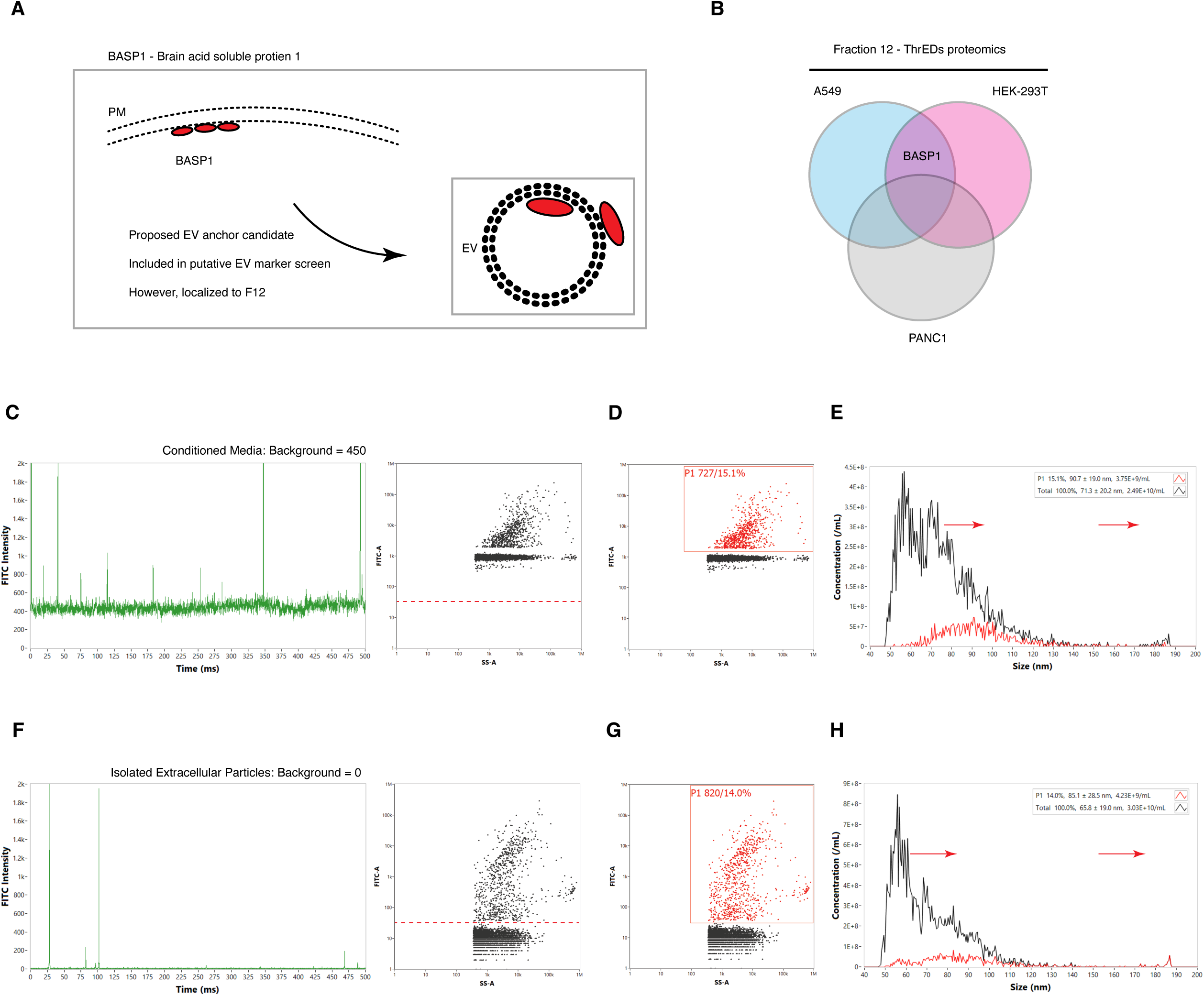
BASP1 is a Thrombospondin Encapsulated Dense Particle protein. (A) Diagram of Brain Acid Soluble Protein 1 (BASP1) a membrane associated cytosolic protein which has typically been described as an extracellular vesicle protein marker (was therefore included in the screen in Figure 1). (B) BASP1 is one of the abundant, common proteins present in ThrED f11-12 samples. (C) Shows the raw FITC readings from the nanoflow cytometer when measuring BASP1-mClover and associated dot plot. The horizontal dotted line represents where a gate would typically be placed resulting in the incorrect gating on BASP1-mClover3 presence in EP samples and a nearly 100% positive FITC population, as has been previously published. (D) Demonstrates where the gate should be placed based on the high background in the sample. (E) Shows the size profile of BASP1 positive and negative extracellular particles with a greater average diameter in positive particles. (F-H) Show the same data from samples isolated using Size Exclusion Chromatography instead of crude conditioned media. (F) Shows lower background when soluble BASP1 proteins are separated from EPs during purification and (G) shows the correct gating. (H) Identifies a similar size profile as in (E). n=3 biological replicates.

**Figure S8.**
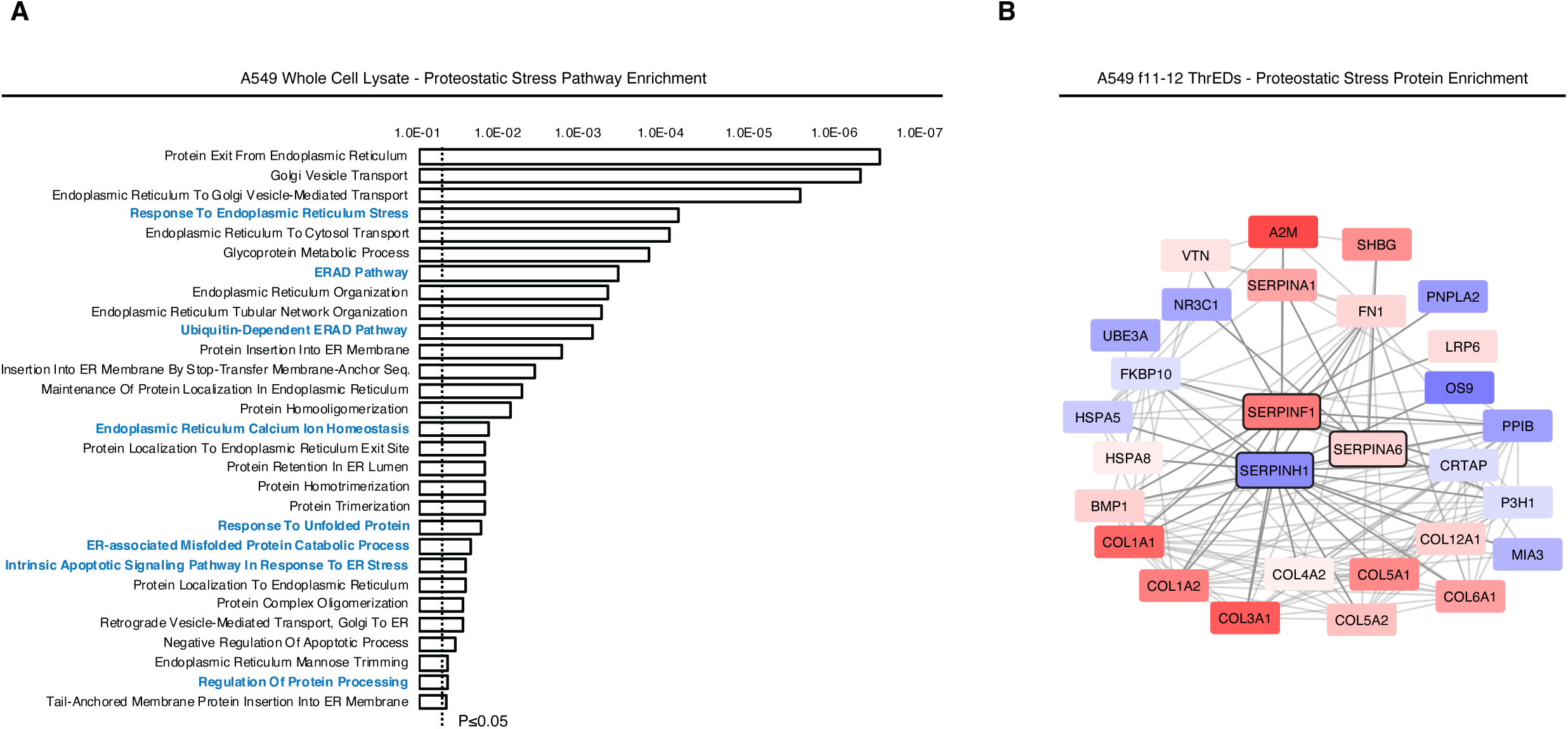
GO term enrichment analysis of proteostatic stress pathways in A549 lysates and ThrEDs. (A) Significant GO terms from proteomics analysis carried out on A549 whole cell lysates at day 4 involved in protein stress response and unfolded protein response and (B) Proteins significantly upregulated in ThrEDs involved in the UPR, cellular stress and ER stress pathways potentially involved in ThrEDs biogenesis and cargo loading. n=3 biological replicates.

**Figure S9.**
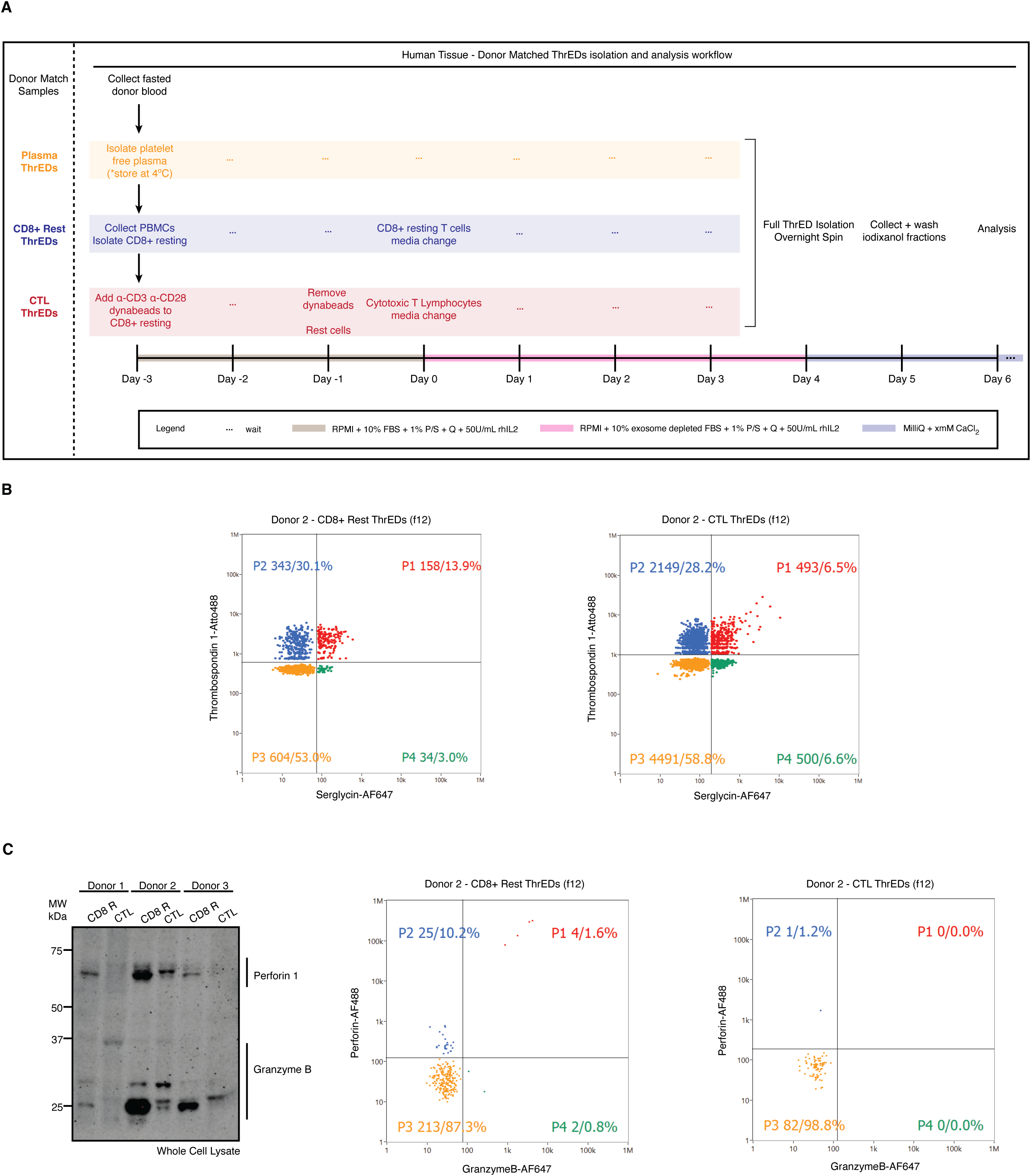
ThrED isolation and characterization from donor matched human tissue. (A) Schematic diagram of protocol timing for ThrEDs in Donor Matched tissues. Fasted blood is collected from donors to generate Platelet Free Plasma, resting CD8+ cells and Cytotoxic T lymphocyte (CTLs). Media used at each step and timing for ThrEDs collection is highlighted. (B) Nanoflow cytometry analysis of f12 ThrEDs from CD8^+^ Resting population and CTLs probing Thrombospondin-1 and Serglycin. (C) Western blot of whole cell lysate from donor resting CD8+ T cells and cytotoxic T lymphocytes probed with Perforin 1 and Granzyme B antibodies to confirm positive cellular expression. (D) Nanoflow cytometry analysis of Donor 2 f12 ThrEDs samples from resting CD8+ cells and CTLs probing Granzyme B and Perforin 1. Data represents n=6 donors. Representative images are shown.

**Figure S10.**
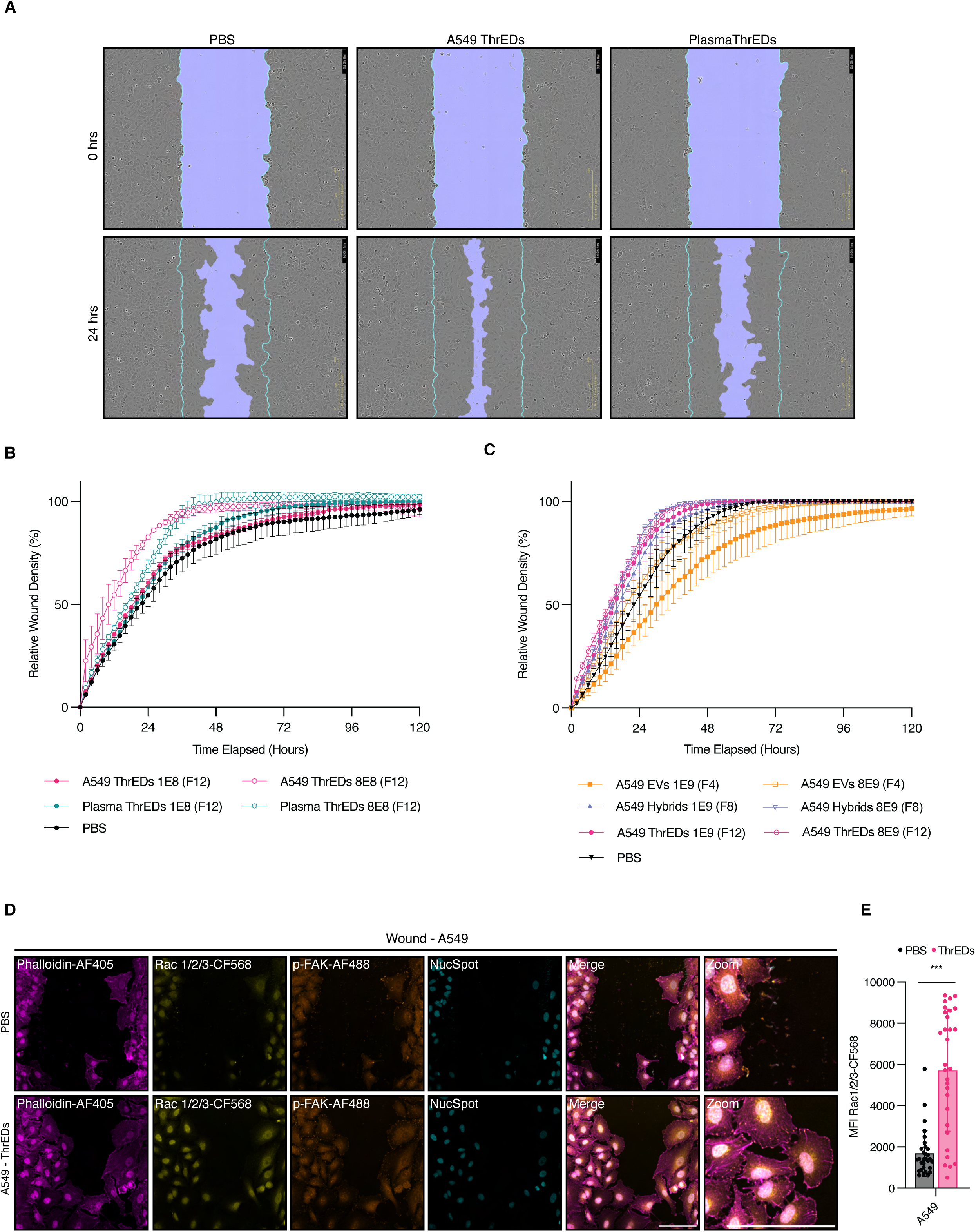
ThrED positive effectors of wound healing in lung adenocarcinoma cells. (A) Representative images of scratch wounds in A549 cells treated with PBS, A549 ThrEDs and Plasma ThrEDs. Initial scratch is shown with a border line (cyan), and the wound area is shown in light purple (the central region). At the 24-hour time point the scratch wound mask fails to recognize certain migrating cells in ThrED treated samples because the cell morphology changes drastically following particle administration to promote migration. (B) Relative wound density of A549 wounds treated with A549 and Plasma ThrEDs from (A) at 2 different doses showing a dose response. (C) Relative wound density of A549 wounds treated with PBS, EVs, Hybrids and ThrEDs at different doses. (D) Confocal imaging of Rac1/2/3 and p-FAK in A549 wounds treated with PBS and ThrEDs at 24 hrs with associated MFI quantification of images (E). Scale bar=100µm. Data represents n=3 biological replicates.

## ACKNOWLEDGEMENTS

We thank E. Kurz, S. Valvo and L. Chen for the assistance with this project; Prof. P. Kubes, Prof. K. Midwood, Prof. M. Coles, Prof. C. Buckley for advice. We acknowledge the generous support of the Kennedy Trust for Rheumatology Research, the Target Discovery Institute Proteomics Facility, IDRM and Carl Zeiss GMBH for the microscopy facilities used in this research including Helena Coker and Kseniya Korobchevskaya, in particular.

## Author Contributions

CCS and MLD conceived the study and designed experiments. CCS performed experiments and analyzed data. CU and CB provided U373 EPs. MW assisted with flow cytometry for Nalm6 particle death assay. AJ provided NK92 SMAPs. RF and IV provided proteomics supervision and data acquisition. AH helped prepare and analyse proteomics data. LW provided myofibroblast cells. DDF and PCD provided Incucyte access. MW provided nanoFCM and ONI access. CS and MLD wrote the manuscript.

